# Genomic loss of the HSP70cA gene in the vertebrate lineage

**DOI:** 10.1101/2022.09.06.506793

**Authors:** Alisha Merchant, Bradly I. Ramirez, Melinda N. Reyes, Dysocheata Van, Marilin Martinez-Colin, Damilola O. Ojo, Esmeralda L. Mazuca, Heidi J. De La O, Abigayle M. Glenn, Claudia G. Lira, Hashimul Ehsan, Ermeng Yu, Gen Kaneko

## Abstract

Metazoan 70 kDa heat shock protein (HSP70) genes have been classified into four lineages: cytosolic A (HSP70cA), cytosolic B (HSP70cB), endoplasmic reticulum (HSP70er), and mitochondria (HSP70m). Because previous studies have identified no HSP70cA genes in vertebrates, we hypothesized that this gene has lost on the evolutionary path to vertebrates. To test this hypothesis, the present study conducted a comprehensive database search followed by phylogenetic and synteny analyses. The HSP70cA gene was present in invertebrates and animals that belong to subphyla of Chordata, Cephalochordata (lancelets) and Tunicata (tunicates). However, the genomes of early vertebrates in the subphylum Craniata (lamprey, hagfish, elephant shark, and coelacanth) contained only HSP70cB genes, suggesting the loss of the HSP70cA gene in the early period of vertebrate evolution. Synteny analysis using available genomic resources indicated that the synteny around the HSP70 genes was generally conserved between tunicates, but it was largely different between tunicates and lamprey. These results suggest the presence of dynamic chromosomal rearrangement in early vertebrates, which possibly caused the loss of the HSP70cA gene in the vertebrate lineage.

## 1. Introduction

The 70-KDa heat shock protein (HSP70) is a family of molecular chaperones present in all three domains of life. Because of the vital importance of HSP70s in maintaining the structure of endogenous proteins, the genes encoding HSP70s have undergone extensive duplication events. For example, *Escherichia coli* possesses three HSP70 genes — *DnaK, HscA*, and *HscC* — that are structurally and functionally distinct to each other (Mayer 2021). In eukaryotes, gene duplication events gave rise to several organelle-specific isoforms. Their subcellular localization of HSP70 changes depending on cellular conditions (Mattos et al. 2022; Mohamad et al. 2022; Velazquez and Lindquist 1984), but the organelle-specific isoforms can be clearly distinguished by taking advantage of phylogenetic analyses. Namely, metazoan HSP70s have been robustly classified into four lineages, cytosolic A (HSP70cA), cytosolic B (HSP70cB), endoplasmic reticulum (ER) (HSP70er), and mitochondria (HSP70m), in Bayesian and maximum likelihood frameworks (Schnebert et al. 2022; Yu et al. 2021), which has been supported by several genome-wide HSP70 screening studies (Grewal et al. 2022; Hasnain and Kaneko; Kaneko 2022; Liu et al. 2022). It should be noted that the metazoan HSP70 family also contains non-canonical members that often show ~30% or less amino acid identities with above HSP70s [e.g., *Homo sapiens* HSP70 4L (NP_055093.2); also see Grewal et al. (2022)]. Considering that *E. coli* DnaK (WP_023278178.1) and *H. sapiens* HSP70-1A (NSPA1A, NP_005336.3) share ~50% amino acid identity, these non-canonical HSP70 genes must have diverged very early in their evolutionary history of eukaryotes.

Among the four canonical HSP70 lineages, the HSP70cAs have shown several unique features. First, HSP70cAs have been found only from invertebrates including Cnidaria, Priapulida, Mollusca, Rotifera, Annelida, and Arthropoda (Yu et al. 2021), suggesting a possible gene loss event during the evolution in the chordate lineage. Second, HSP70cAs have a characteristic serine residue in the ATPase domain with a few exceptions (Drosopoulou et al. 2009; Grewal et al. 2022; Kourtidis et al. 2006; Yu et al. 2021), although the function of this residue remains to be elucidated. Third, the HSP70 family members with a relatively constitutive expression pattern, which have been traditionally called the 70 kDa heat shock cognates (HSC70), exclusively belong to the HSP70cB lineage (Yu et al. 2021). Accordingly, all known HSP70cA genes are either demonstrated to be stress-inducible or are not known for their expression patterns. These features make the HSP70cA gene an attractive target of molecular evolutionary research.

In the present study, given the apparent absence of the HSP70cA genes in human and other vertebrates in our previous studies, we hypothesized that the HSP70cA gene has lost on the evolutionary path to vertebrates. To test this hypothesis, we conducted a large-scale database screening followed by comparative genome analysis.

## 2. Materials and Methods

### 2.1. NCBI database screening

The NCBI nucleotide database was screened for metazoan HSP70s on October 26, 2021, using several different keywords and filters (Supplementary Table S1). Screened sequences were tested for the presence of the HSP70cA-specific serine residue by Clustal Omega alignment with two partial HSP70cA sequences: ILTIDEGSLFEVRSTAGD from *Drosophila melanogaster* HSP70Aa (HSP70cA1, AAG26887.1) and VLAIDEGSIFEVKATAGD from *Aplysia californica* HSP70cA1 (XP_005103834.1). The screened sequences were further analyzed with in-house zsh and R scripts using following functions. Duplicated sequences were removed by the seqkit rmdup using sequence as an identifier. Clustal W (version 1.2.4) and was used for the pairwise comparison of the screened sequences with the 20 HSP70cA and 46 HSP70cB amino acid sequences (Supplementary Table S2). Pairwise alignment scores were extracted, and the average scores were calculated. R (version 4.1.2) was used to make the histogram and heatmap of the alignment scores.

### 2.2. Phylogenetic and synteny analyses

Sequence alignment was performed using Clustal Omega (clustalo v1.2.4) (Sievers et al. 2011), M-Coffee (Wallace et al. 2006), and/or MUSCLE v3.8.1551. Best-fit models were determined for each alignment by ProtTest-3.4.2 (Darriba et al. 2011) using the Bayesian information criterion. Bayesian trees were constructed using MrBayes 3.2.7a (Ronquist et al. 2012) with 500,000 – 1,000,000 generations. Every 10th tree was sampled, and burn-in was set to 25%. Maximum-likelihood trees were constructed using MEGA11 with 1000 bootstrap replications. Phylogenetic trees were visualized with FigTree (v1.4.4) or MEGA11 using the mitochondrial lineage as the root. Synteny analysis was performed using the DECIPHER package of R.

### 2.3. Genome-wide screening

The following datasets were used for the genome-wide screening of HSP70: vase tunicate *Ciona intestinalis* Ensembl peptide sequences release 105 (17,302 sequences from assembly GCA_000224145.1), Pacific transparent sea squirt *Ciona savignyi* Ensembl peptide sequences release 105 (20,155 sequences from assembly CSAV 2.0), Florida lancelet *Branchiostoma floridae* Emsembl Metazoa peptide sequences release 52 (36,011 sequences from assembly GCA_900088365.1), sea lamprey *Petromyzon marinus* Ensembl peptide sequences release 107.7 (assembly Pmarinus_7.0), inshore hagfish *Eptatretus burgeri* peptide sequences release 107.32 (assembly GCA_900186335.2), elephant shark *Callorhinchus milii* peptide sequences release 6.1.3 (assembly GCA_000165045.2), and coelacanth *Latimeria chalumnae* peptide release 107.1 (GCA_000225785.1). Local BLASTP was performed using the amino acid sequence of human HSP70cB1 (HSPA1A, NP_005336.3) as a query.

## 3. Results and Discussion

### Gene screening

The NCBI nucleotide database search using various keywords related to HSP70s (Supplementary Table S1) resulted in the identification of 6,572 sequences (Fig. 1A). After manually adding 20 HSP70cA and 46 HSP70cB sequences characterized in our previous study (Supplementary Table S2) (Yu et al. 2021), 2,804 duplicated sequences were removed. In order to screen HSP70s similar to known HSP70cAs rather than HSP70cBs, the remaining 3,834 sequences were used for the pairwise comparison against the 20 HSP70cA and 66 HSP70cB sequences that have been previously characterized (Supplementary Table S2). The pairwise alignment score showed a wide distribution from 3.45 to 100 with two three peaks around 10, 30, and 75 (Fig. 1B). Closer data inspection revealed that the 3,834 sequences contained non-canonical HSP70 family members called HSP12-14, which generally showed the pairwise alignment scores of about 10 to 30. The screened sequences also contained proteins that are not homologous to HSP70s, such as HSP70-interacting proteins. Given that *Saccharomyces cerevisiae* SSA1 (YAL005C) and *Drosophila melanogaster* HSP70cA1 (AAG26887.1) showed the pairwise alignment score of 71 (i.e., 71% amino acid identity), we set the threshold at 60 to narrow down our search for HSP70cA genes to 2,049 genes.

**Fig. 1.**
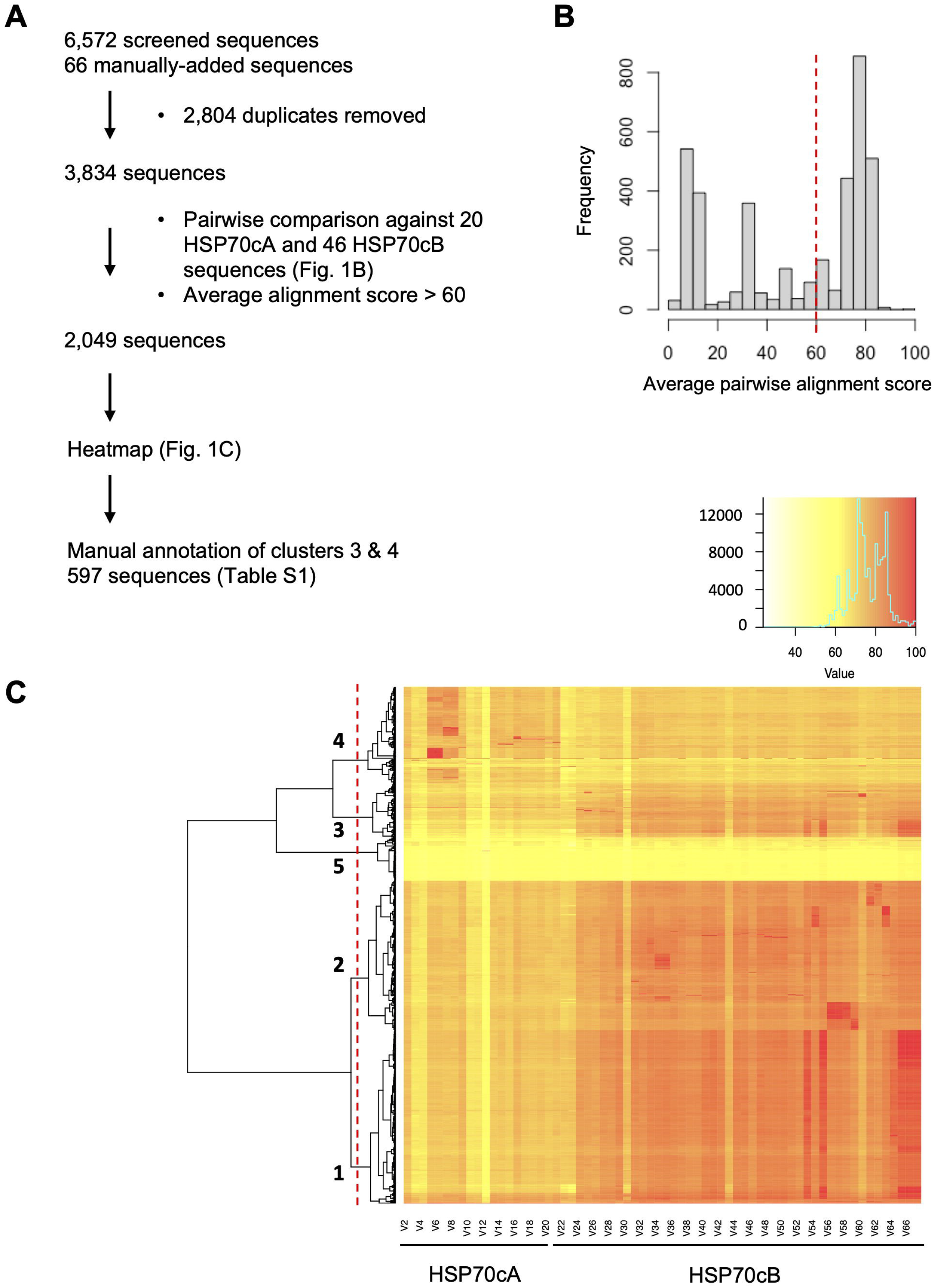
NCBI database screening for the discovery of HSP70cAs. (A) Screening strategy. (B) Distribution of average pairwise alignment scores between each screened HSP70 and 66 HSP70cA/HSP70cB proteins listed in Supplementary Table S2. The red broken line indicates the cutoff threshold (alignment score = 60). (C) Clustering analysis of the pairwise alignment score. Pairwise alignment scores were calculated between each of the screened HSP70 (Y axis) and the 66 HSP70cA/HSP70cB proteins (Supplementary Table S2). Red broken line indicates the clustering threshold, with which screened HSP70 proteins in the cluster 4 (383 proteins) show higher pairwise alignment scores to HSP70cAs than to HSP70cBs.

The 2,049 sequences were separated into clusters 1 to 5 containing 688, 592, 214, 383, and 172 sequences, respectively, depending on the alignment scores to the 20 HSP70cA and 46 HSP70cB proteins (Fig. 1C). The 383 HSP70-like proteins in the cluster 4 showed higher alignment scores to the 20 HSP70cA proteins than to 46 HSP70cB proteins, constituting the pool of possible HSP70cA proteins in the NCBI nucleotide database.

### Phylum distribution

The 383 HSP70s in the cluster 4 were manually examined for the DNA accession numbers, protein accession numbers, and the presence of the HSP70cA-specific serine residue, together with the scientific name, phylum, and class of the species (Supplementary Table S3). We also annotated 214 sequences in the cluster 3 as a control group because sequences in the cluster 3 generally showed higher similarities to HSP70cBs than to HSP70cAs (Fig. 1C). During the manual annotation, we removed 4 and 3 sequences from clusters 3 and 4, respectively, because of their low quality and/or short length that did not cover the serine residue in the ATPase domain.

Fig. 2 shows the phylum distribution of the sequences in clusters 3 (210 HSP70s) and 4 (380 HSP70s). The cluster 3 contained many chordate HSP70s without the HSP70cA-specific serine residue. The cluster 3 also contained HSP70s from various phyla, but only a few of them had the serine residue specific to the HSP70cA lineage. On the other hand, the cluster 4 contained many arthropod HSP70s with the serine residue (putative HSP70cAs). Compared to the HSP70s from Arthropoda and Mollusca, there were significantly lower number of Chordata proteins in the cluster 4 with five sequences that did not have the serine residue. Overall, our NCBI database search combined with the cluster analysis highlighted the tendency that the putative HSP70cAs are dominant in Arthropoda and Mollusca, but minor in Echinodermata, Hemichordata, and Chordata. These results prompted us to perform a detailed sequence inspection of HSP70s in the clusters 3 and 4 with a special attention to HSP70 family members from Echinodermata, Hemichordata, and Chordata.

**Fig. 2.**
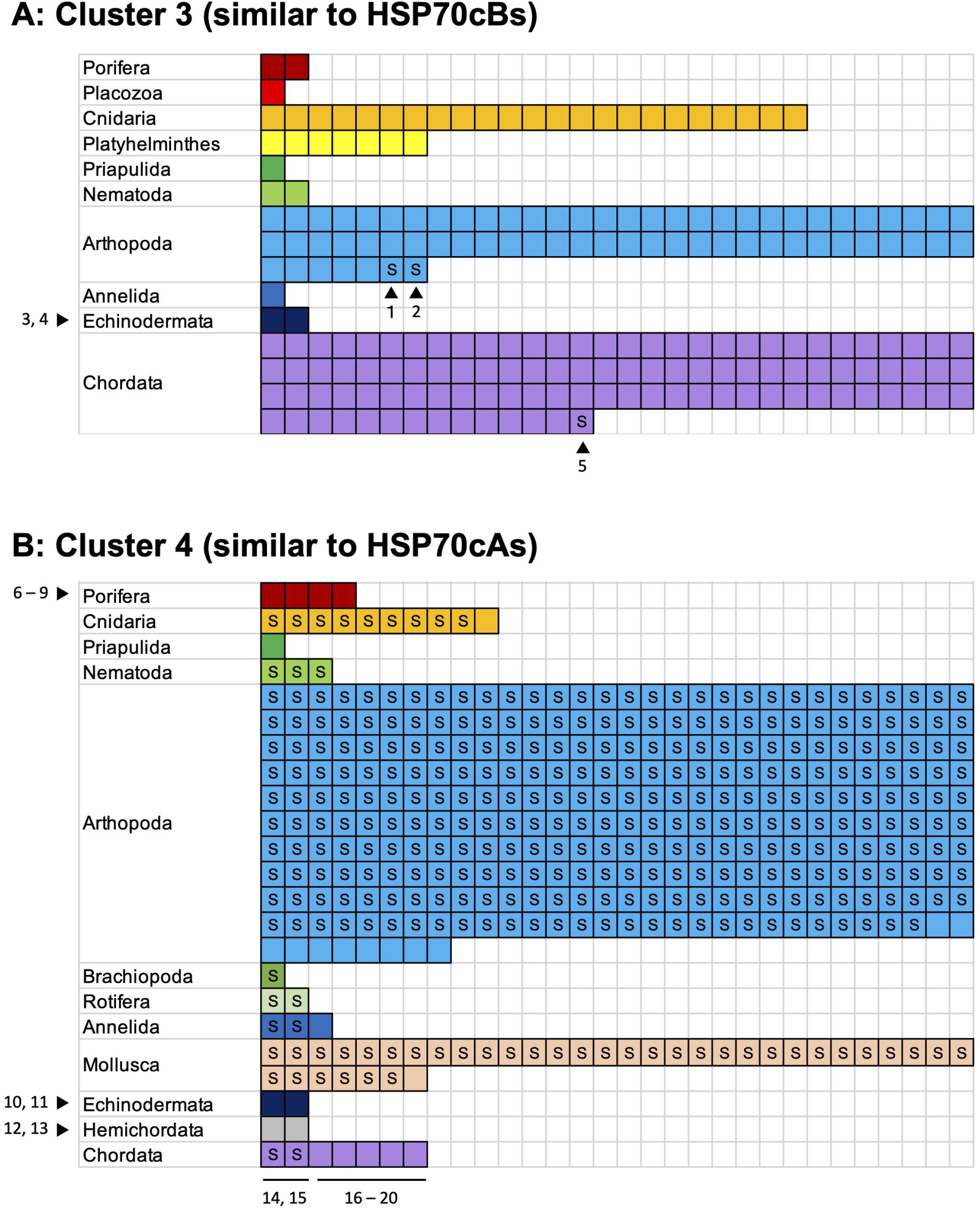
Phylum distribution of the screened HSP70 genes. The number of boxes indicate the number of HSP70s screened from the phylum. “S” in each box indicates that the HSP70 protein contained the serine residue specific to the HSP70cA lineage. Numbers (1 to 20) are corresponding to those in Fig. 3.

### Phylogenetic analysis

We constructed Bayesian and ML phylogenetic trees including the 20 HSP70 sequences that showed irregular distribution patterns in Fig. 2 (numbered 1 – 20). To make sure that the trees contain the four lineages of canonical HSP70s (cytosolic A, cytosolic B, ER, and mitochondria), 26 HSP70 sequences annotated in previous studies were also included (Supplementary Table 4). All nodes were well resolved with a high posterior probability. The topology of these phylogenetic trees generally supported findings of previous studies (Grewal et al. 2022; Yu et al. 2021). First, the four lineages were clearly separated, and yeast HSP70s were basal to metazoan HSP70s in each lineage, confirming the ancient separation of the four lineages. Second, the cytosolic lineage was further separated to A and B, and the extra serine residue was absent in the lineage B. The serine residue was present in only in HSP70cA sequences, although it was substituted in Porifera, Echinodermata, and Hemichordata HSP70cAs. In addition, these results were consistent in the six phylogenetic trees constructed in this study (Bayesian and ML × three alignments: Clustal Omega, MUSCLE, and M-Coffee; data now shown), indicating that the results are robust to changing analytical methods and addition of low-quality sequences (see below).

HSP70s with the irregular distribution patterns in Fig. 2 were assigned to either the HSP70cA or HSP70cB lineage. Arthropoda HSP70s with the extra serine residue (Nos. 1 and 2 in Fig. 2 from *Pentalonia nigronervosa* and *Sipha flava*) turned out to be HSP70cA, although they showed higher amino acid alignment scores to HSP70cBs and thus assigned to the cluster 3. These were partial HSP70 sequences, which possibly affected the result of the classification based on the amino acid alignment scores. Porifera HSP70s (Nos. 6 – 9) in the cluster 3 were assigned to both HSP70cA (Nos. 6 and 9) and HSP70cB (Nos. 7 and 8) clusters. This inconsistency between Figs. 2 and 3 is probably attributed to the low amino acid identity of Porifera HSP70s to other metazoan HSP70s. The four Echinodermata HSP70s belonged to HSP70cA (Nos. 10, 11) and HSP70cB (Nos. 3, 4) clusters, which was consistent between Figs. 2 and 3. Hemichordata HSP70s (Nos. 12 and 13) were in the HSP70cA cluster, although they did not have the extra serine residue. Chordata HSP70s with the serine residue (Nos. 14 and 15) belonged to the HSP70cA cluster, whereas those without the serine residue (Nos. 16 – 20) belonged to HSP70cB. The Chordata HSP70cBs were either partial (No. 16) or tagged as “low quality protein” (Nos. 17 – 20) (Supplementary Table 4), which may have caused the inconsistency between Figs. 2 and 3. Importantly, the two Chordata HSP70cAs were from tunicates, a subgroup of Chordata, and all vertebrate HSP70 belonged to the lineage B.

**Fig. 3.**
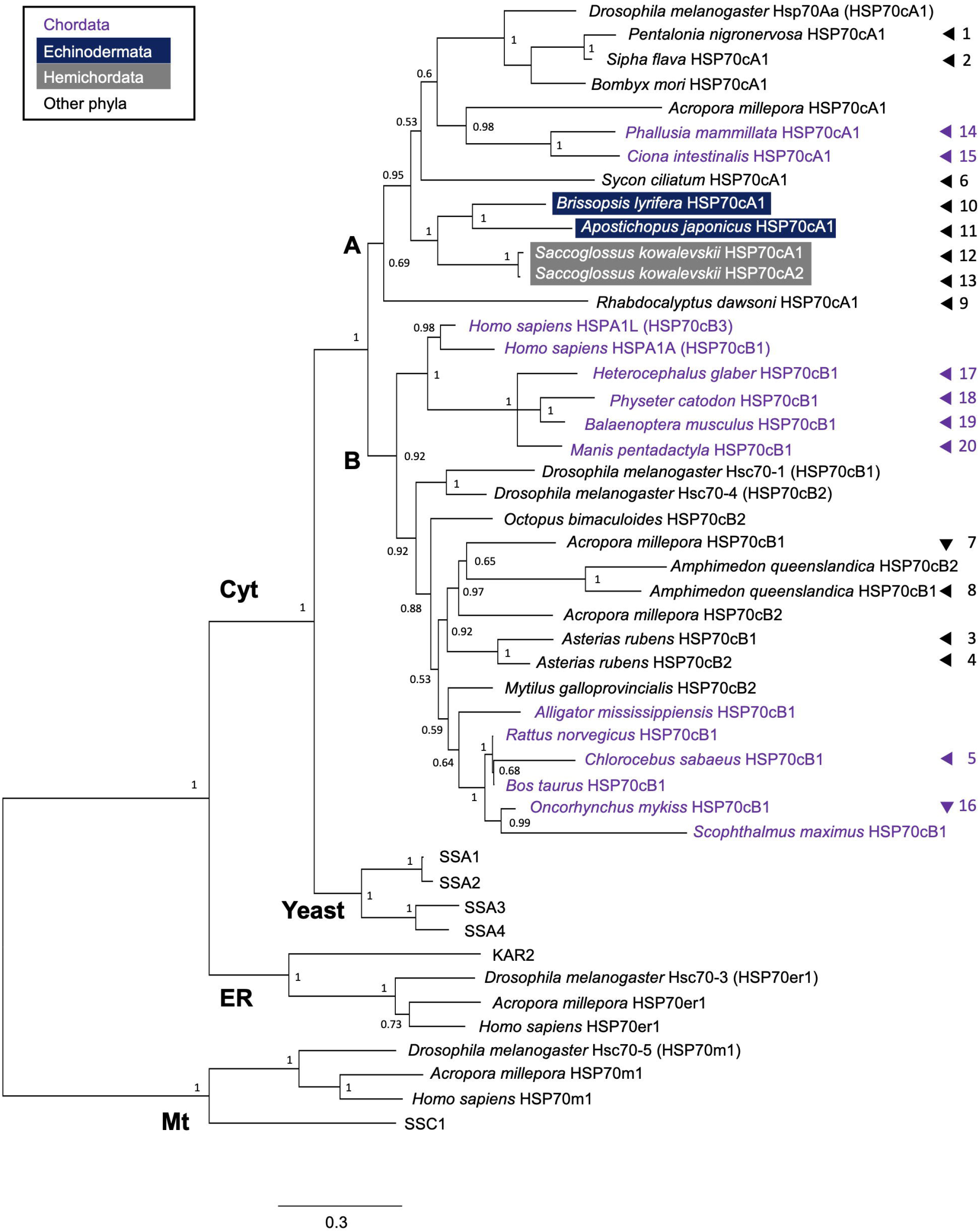
Bayesian consensus tree of HSP70 family members. HSP70 sequences that showed irregular characteristics in Fig. 2 were included. Numbers on the right margin correspond to those in Fig. 2, whereas numbers on the branches are the posterior probability support for each node. The detailed sequence information including accession numbers are shown in Table 1. The M-Coffee program was used for the alignment with default parameters, and the phylogenetic tree was constructed using the LG+G substitution model and 1,000,000 generations.

Overall, the phylogenetic analysis enabled us to further refine the annotation process, clarifying the lineages of some low-quality HSP70s. This analysis demonstrated the presence of HSP70cA in Echinodermata, Hemichordata, and Chordata (tunicates). The apparent absence of HSP70cA in human thus can be explained by the gene loss within the phylum Chordata, which took place after the split of vertebrates from the common ancestor with tunicates.

### Chordata analysis

The phylum Chordata contains three subphyla, Cephalochordata (lancelets), Craniata (vertebrates), and Tunicata (tunicates) in the NCBI Taxonomy database. Most studies agree with this classification, although there are some variations in the classification level (Irie et al. 2018; Satoh et al. 2014). Molecular evidence suggests that Cephalochordata diverged from the other two groups more than 520 million years ago. Therefore, we screened HSP70 genes in Cephalochordata and Tunicata genomes to examine whether they contain HSP70cA genes.

We first conducted a local BLASTP search against the collection of peptide sequences deduced from the genome of European lancelet *Branchiostoma floridae* using human HSP70cB1 sequence as a query. Four sequences showed BLAST scores more than 500 (Table 1). The fifth top hit with the BLAST score of 340 (BL04064) encoded *B. floridae* HSP70-14, suggesting that this species has only four canonical HSP70 genes. These sequences were subjected to the phylogenetic annotation together with Tunicata HSP70 sequences identified below.

**Table 1.** Cephalochordata and Tinicata HSP70 genes used in the phylogenetic analysis

| Species | Ensembl Protein ID | BLAST score* | Ensembl Gene ID | Published name | Proposed name | Note, variant, and/or location |
| --- | --- | --- | --- | --- | --- | --- |
| *Branchiostoma lanceolatum* | - | 839 | BL10970 | - | HSP70cB1 | Sc0000030:2316840-2321566 |
| *Branchiostoma lanceolatum* | - | 782 | BL18316 | - | HSP70er1 | Sc0000198:367558-389400 |
| *Branchiostoma lanceolatum* | - | 637 | BL21182 | - | HSP70cA1 | Sc0000038:1645245-1647044 |
| *Branchiostoma lanceolatum* | - | 575 | BL04064 | - | HSP70m1 | Sc0000207:169399-182801 |
| *Ciona intestinalis* | ENSCINP00000019652.3 | 1028 | ENSCING00000009665 | Ci-HSPA1/6/7-like | HSP70cB1 | HT000062.1:524353-528965 |
| *Ciona intestinalis* | ENSCINP00000011757.3 | 915 | ENSCING00000005686.3 | Ci-HSPA2/8 | HSP70cA1 | Same as NP_001029006.1 in Table 1, Scaffold HT000129.1: 76,239-78,122 |
| *Ciona intestinalis* | ENSCINP00000019494.3 | 792 | ENSCING00000009590.3 | Ci-HSPA5b | HSP70er2 | Chr 9:5050624-5054713 |
| *Ciona intestinalis* | ENSCINP00000019205.3 | 718 | ENSCING00000009445.3 | Ci-HSPA5a | HSP70er1 | Chr 9:4838560-4843040 |
| *Ciona intestinalis* | ENSCINP00000033187.1 | 559 | ENSCING00000019791.1 | Ci-HSPA9B | HSP70m1 | Chr 3:5760246-5762192 |
| *Ciona savignyi* | ENSCSAVP00000010977.1 | 1056 | ENSCSAVG00000006429.1 | Hsp70-2** | HSP70er1 | Variant ENSCSAVP00000010978.1, reftig_140:663164-666859 |
| *Ciona savignyi* | ENSCSAVP00000007990.1 | 1025 | ENSCSAVG00000004764.1 | Hsp70-3** | HSP70cB1 | Variant ENSCSAVP00000007989.1, reftig_117:237661-240908 |
| *Ciona savignyi* | ENSCSAVP00000015814.1 | 956 | ENSCSAVG00000009296.1 | Hsp70-4** | HSP70cA1 | Variant ENSCSAVP00000014698.1, reftig_11:1,397,712-1,399,655 |
| *Ciona savignyi* | ENSCSAVP00000014697.1 | 938 | ENSCSAVG00000008588.1 | Hsp70-5** | HSP70cB2 | reftig_49:1564826-1572778 |
| *Ciona savignyi* | ENSCSAVP00000017229.1 | 903 | ENSCSAVG00000010133.1 | Hsp70-6** | HSP70cB3 | reftig_58:2768858-2774321 |

\*Against human HSP70cB1
\*\* Huang et al., 2018

In Tunicata, several HSP70 genes have been identified from vase tunicate *Ciona intestinalis* (Wada et al. 2006) and Pacific transparent sea squirt *C. savignyi* (Huang et al. 2018). However, we conducted the BLAST screening again because of the updates in the genomic sequence and unavailability of some information in these previous studies. In the *C. intestinalis* genome, amino acid sequence of five HSP70 genes showed a high BLAST score with human HSP70cB1 (HSPA1A) (Table 1). These genes likely correspond to the five canonical HSP70s reported previously (Wada et al. 2006), although the Ensembl database number was not provided in the previous study. As for *C. savignyi*, Huang et al. (2018) identified eight HSP70 genes, two of which encoded non-canonical HSP70s. Another HSP70 gene (ENSCSAVG00000009801) identified by Huang et al. (2018) did not exist in the latest Ensembl peptide data since this gene was annotated as a pseudogene. The remaining five HSP70 genes were reproducibly screened in this study with high BLAST scores (Table 1). The presence of only about five canonical HSP70 genes in Cephalochordata and Tunicata genomes reinforced the previously reported idea that the HSP70 family increased the number and diversity in the lineage to human after the separation from the *Ciona* lineage (Wada et al. 2006).

The phylogenetic analysis successfully classified these Cephalochordata and Tunicata HSP70s into the four lineages (Fig. 4A), which was supported by the signature sequences (Fig. 4B) as well as by the presence and absence of serine residue (Fig. 4C). These results demonstrated the presence of the HSP70cA genes in subphyla Cephalochordata and Tunicata, implying that the HSP70cA gene was lost in the Craniata lineage after the separation from the Tunicata lineage. It is noted that we did not find the HSP70m gene in the *C. savignyi* genome, although the gene was present in both *B. floridae* and *C. intestinalis* genomes. This may be attributed to the incomplete sequencing of the *C. savignyi* genome; the number of predicted genes in the *C. savignyi* genome was only 11,616, although the *C. intestinalis* genome contained 16,671 predicted coding genes.

**Fig. 4.**
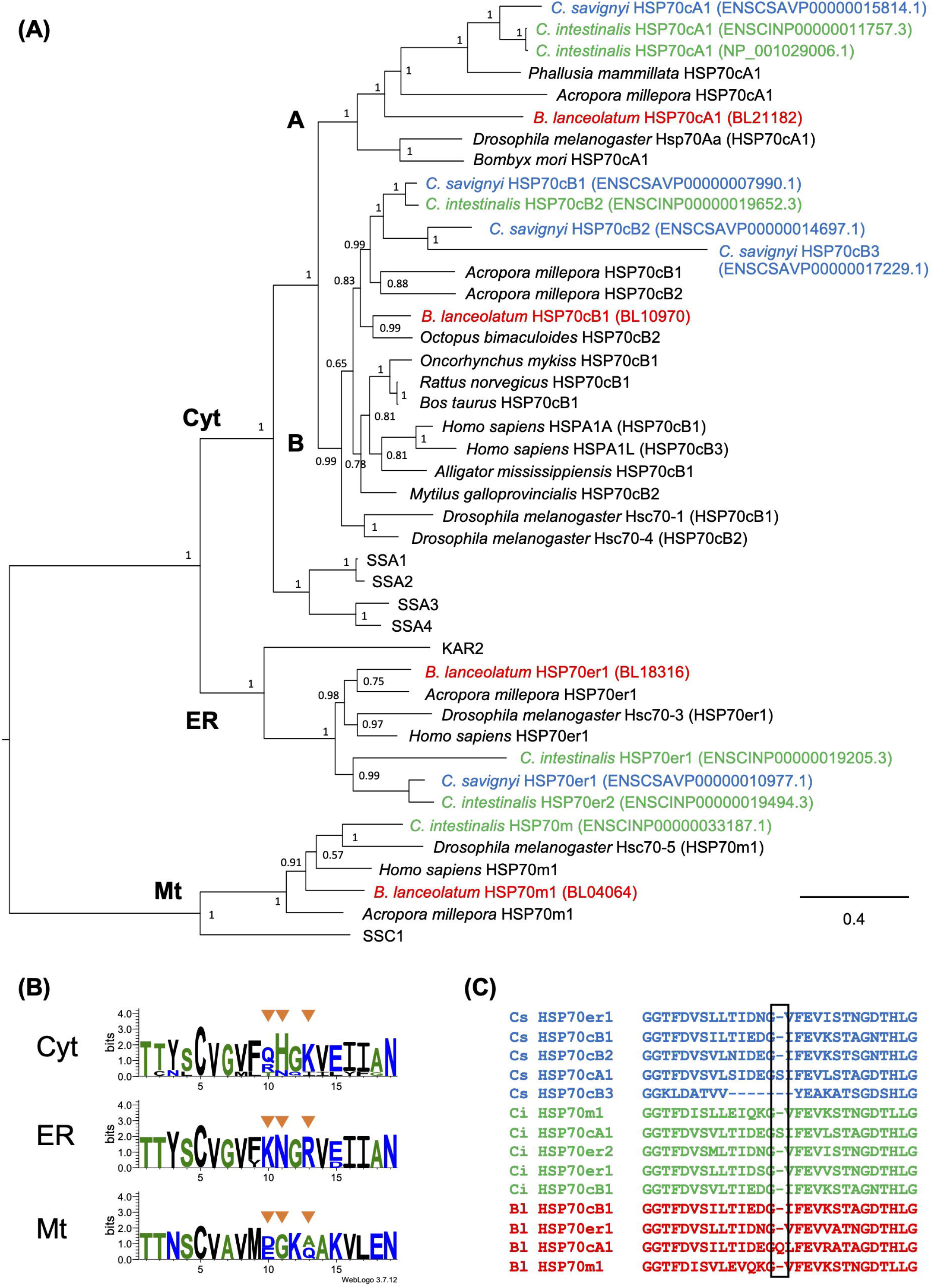
Phylogenetic annotation of Cephalochordata and Tunicata HSP70s. (A) Bayesian consensus tree of canonical HSP70 family members identified by genome-wide screening. Numbers on the branches indicate the posterior probability support. The detailed sequence information including accession numbers are shown in Table 1. The M-Coffee program was used for the alignment with default parameters, and the phylogenetic tree was constructed using the LG+G substitution model and 1,000,000 generations. (B) Sequence logo created from the 14 Cephalochordata and Tunicata HSP70s in the phylogenetic tree. The *B. floridae* HSP70cB1 was removed from the alignment in this analysis because it was a partial sequence missing the aligned area. (C) Presence and absence of the HSP70cA-specific serine residue in Cephalochordata and Tunicata HSP70 sequences.

### Vertebrata analysis

The Ensembl database “Other vertebrates” category contained genomes from four early vertebrates including the lamprey *P. marinus*, hagfish *E. burgeri*, elephant shark *C. milii*, and coelacanth *L. chalumnae*. To examine whether the HSP70cA gene is present in these genomes, we conducted the genome-wide screening for these four species (Table 2). BLASTP search against the lamprey *P. marinus* genome found only two HSP70-like genes, possibly due to the low coverage. The hagfish *E. burgeri* and coelacanth genomes contained 4 and 10 canonical HSP70 genes, respectively. Surprisingly, the genome of elephant shark *C. milii* contained 30 canonical HSP70 genes, likely reflecting the whole-genome duplication event. Interestingly, the phylogenetic analysis indicated that none of these vertebrate HSP70 genes belonged to the HSP70cA cluster (Fig. 6). We therefore concluded that the HSP70cA gene was lost during early vertebrate evolution.

**Fig. 5.**
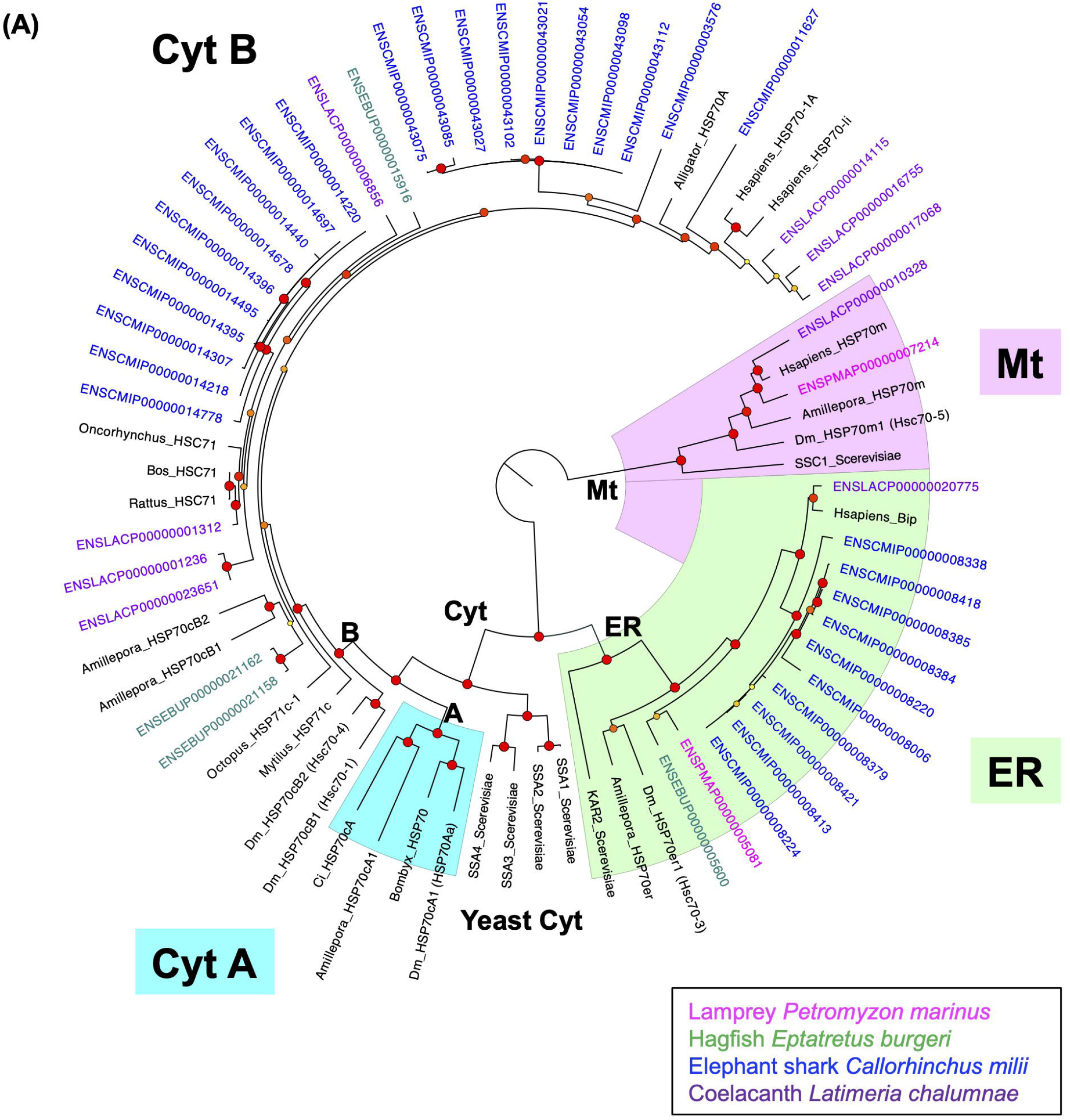
Bayesian consensus tree of HSP70 family members. HSP70 sequences from lamprey, hagfish, elephant shark, and coelacanth were included. Numbers on the branches indicate the posterior probability support. The detailed sequence information including accession numbers are shown in Table 1. The M-Coffee program was used for the alignment with default parameters, and the phylogenetic tree was constructed using the LG+G substitution model and 1,000,000 generations.

**Fig. 6.**
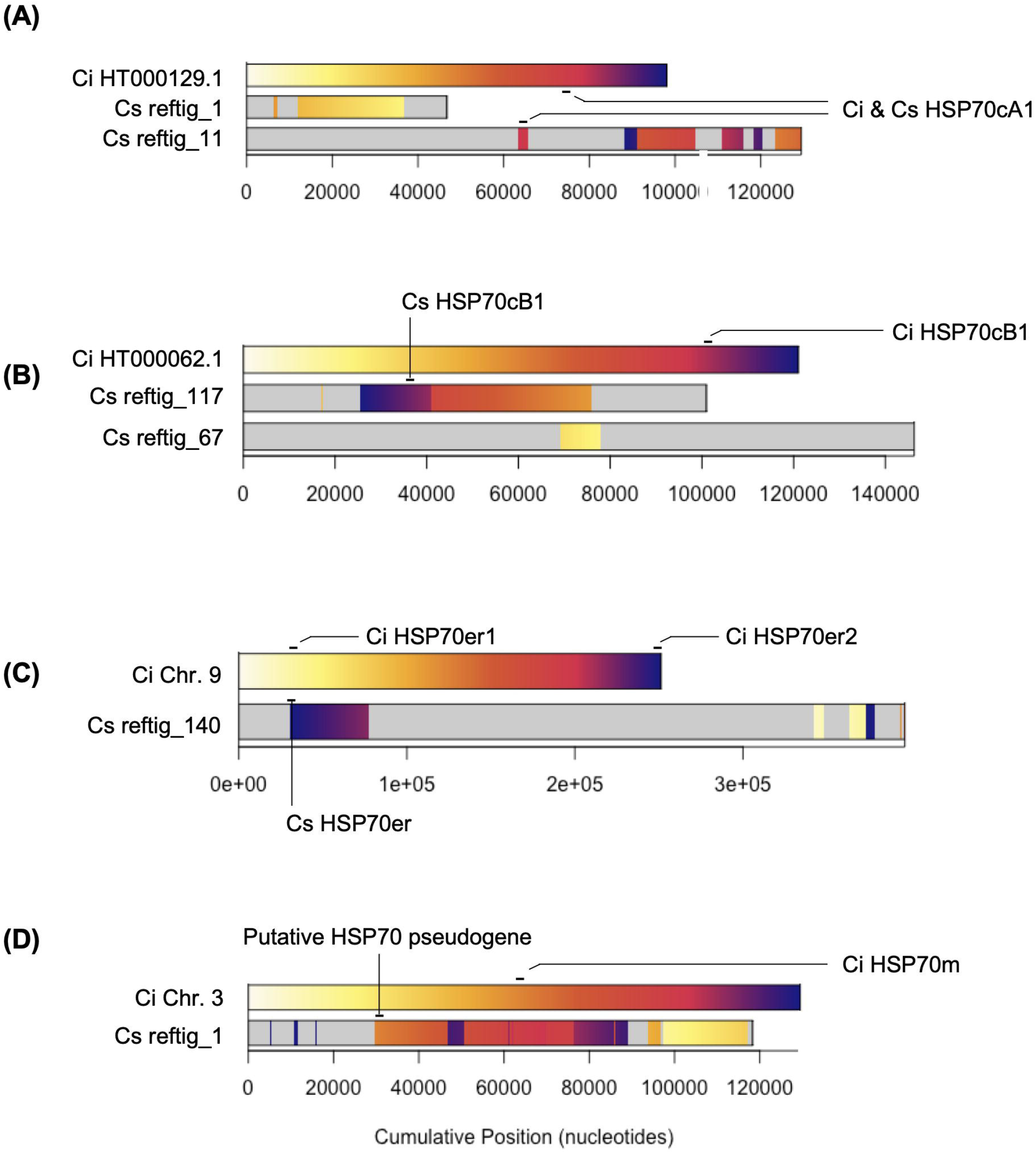
Synteny analysis between genomic regions containing the HSP70 genes. Approximate positions of HSP70 genes are indicated. The same color indicates aligned region. (A) Ci HT000129.1 = *C. intestinalis* KH:HT000129.1:1:98070:1, Cs reftig 1 = *C. savignyi* reftig_1:3133601:3180338:1, Cs reftig 11 = *C. savignyi* reftig_11:1333969:1463440:1. (B) Ci HT000062.1 = *C. intestinalis* KH:HT000062.1:422312:543332:1, Cs reftig_117 = *C. savignyi* reftig_117:198714:299703:1, Cs reftig_67 = *C. savignyi* reftig_67:474648:620967:1. (C) Ci Chr. 9 = *C. intestinalis* KH:9:4803533:5054713:1, Cs reftig_140 = reftig_140:632968:1029275:1. (D) Ci Chr. 3 = *C. intestinalis* KH:3:5696489:5825960:1, Cs reftig_1 = reftig_1:1969128:2087431:1.

**Table 2.**
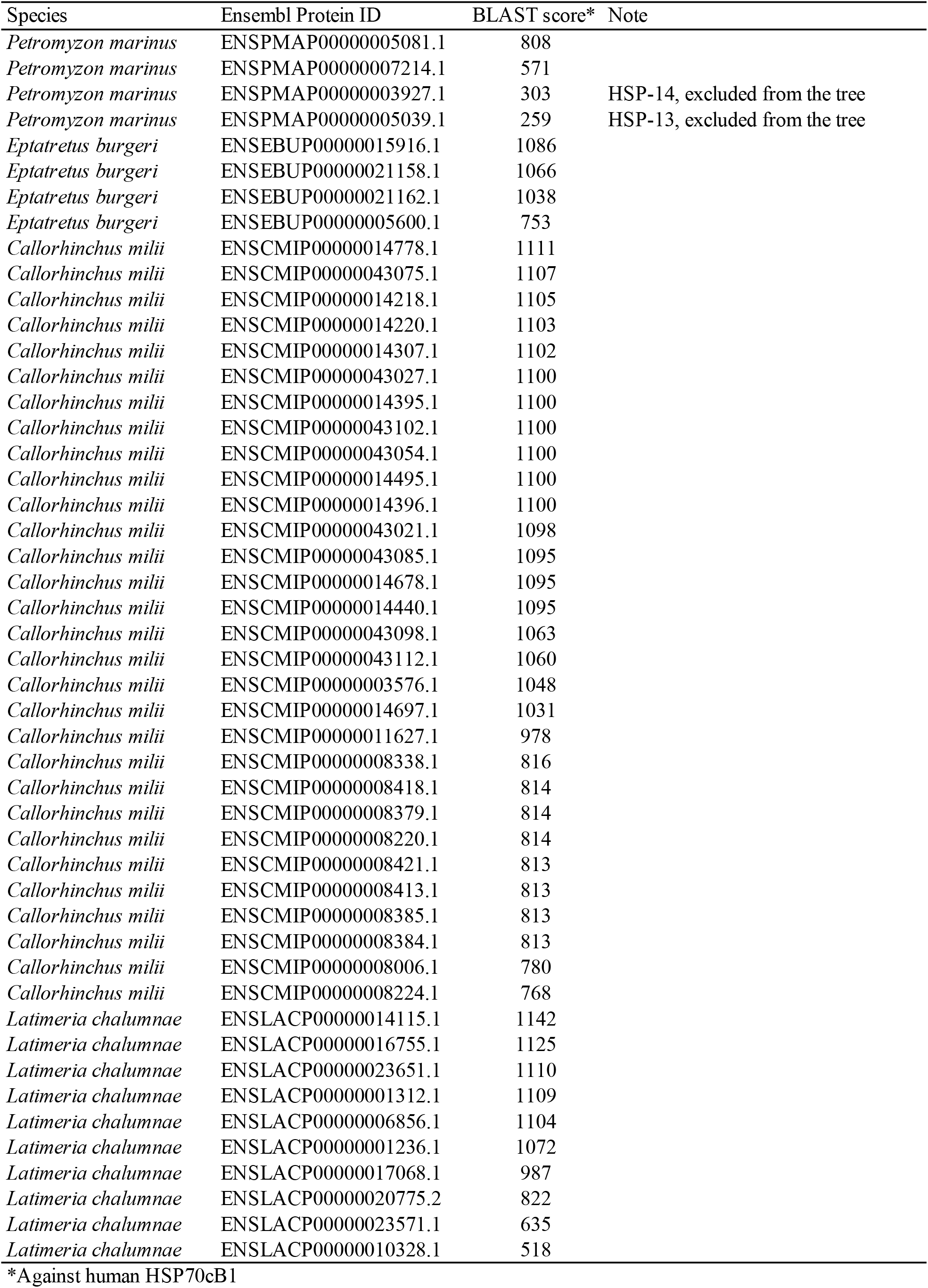
Craniata HSP70 genes used in the phylogenetic analysis

### Synteny analysis

Lastly, we compared synteny of HSP70cA and HSP70cB genes in the genomes of *C. intestinalis, C. savignyi*, and *B. floridae* genomes to gain insights into how the HSP70cA gene was lost from vertebrate genomes. In the Ensembl database, *C. intestinalis* HSPcA1 was located on the HT000129.1, which was aligned mainly to contigs reftig_1 and reftig_11 of *C. savignyi* genome (Fig. 6A). While the *C. intestinalis* HT000129.1 corresponded to the two *C. savignyi* contigs, due to either chromosome rearrangement or incomplete assembly, the synteny around the tunicate HSP70cA1 gene was generally conserved. Similarly, the *C. intestinalis* HT000062.1 containing the HSP70cB1 gene corresponded mainly to reftig_117 and reftig_67 of the *C. savignyi* genome (Fig. 6B), and the synteny in these regions were generally conserved, indicating that the HSP70cB1 genes are orthologous in these two species. The two *C. intestinalis* HSP70er genes (Table 1 and Fig. 4) were located tandemly on the chromosome 9 (Fig. 6C), whereas the *C. savignyi* genome contained only one HSP70er ortholog in the corresponding region (reftig_140), indicating a gene duplication event in *C. intestinalis* or a loss of the HSP70er gene in *C. savignyi. C. intestinalis* chromosome 3 containing HSP70m was aligned with *C. savignyi* reftig_1, but there was no *C. savignyi* HSP70m ortholog in this region in line with the absence of the HSP70m gene in the *C. savignyi* genome (Fig. 4). However, detailed sequence inspection identified a putative HSP70m pseudogene of *C. savignyi* in this region (Fig. 6D, ENSCSAVG00000009801, reftig_1: 1,998,692– 2,000,750). The deduced amino acid sequence of this gene showed ~93% amino acid identity to *C. intestinalis* HSP70m, but N- and C-terminal sequences seemed missing, and there was no available EST data that supported the transcription of this region. Taken together, the synteny analysis demonstrated that chromosomal regions containing the HSP70 genes were generally conserved between *C. intestinalis* and *C. savignyi*, but there were some duplication or gene loss even between the two closely related tunicate species in the same genus.

We subsequently aligned the *C. intestinalis* genomic regions containing HSP70cA, HSP70cB, HSP70er1, and HSP70m genes with the *P. marinus* genome. When small regions containing 5 kb upstream and downstream of *C. intestinalis* HSP70cA, HSP70cB, HSP70er1, HSP70er2 and HSP70m genes were aligned to the *P. marinus* genome, 0.25%, 7.9%, 3.1%, 5.1%, and 0.25% of the regions were mapped, respectively (Fig. 7). To the *C. savignyi* genome, 22%, 23%, 13%, 17%, 25% of these regions were mapped, respectively. When large regions in Fig. 6A – 6D (about 100 kb around the HSP70 genes) were aligned to the *P. marinus* genome, 0.21%, 1.1%, 0.41%, and 1 % of these regions were aligned, respectively (Fig. 8). To the *C. intestinalis* genome, 4.7%, 4.9%, 7.9%, and 7.7 % of these regions were mapped, respectively. The alignment patterns were consistent with the difference; there were several continuously aligned regions between the *C. intestinalis* genomic regions and *C. savignyi* genome, but alignment with the *P. marinus* genome was quite intermittent, indicating that the synteny around HSP70 genes was generally not well conserved between *C. intestinalis* and *P. marinus*. However, importantly, regions around the *C. intestinalis* HSP70cA1 gene consistently showed the lowest mapping rate to the *P. marinus* genome.

**Fig. 7.**
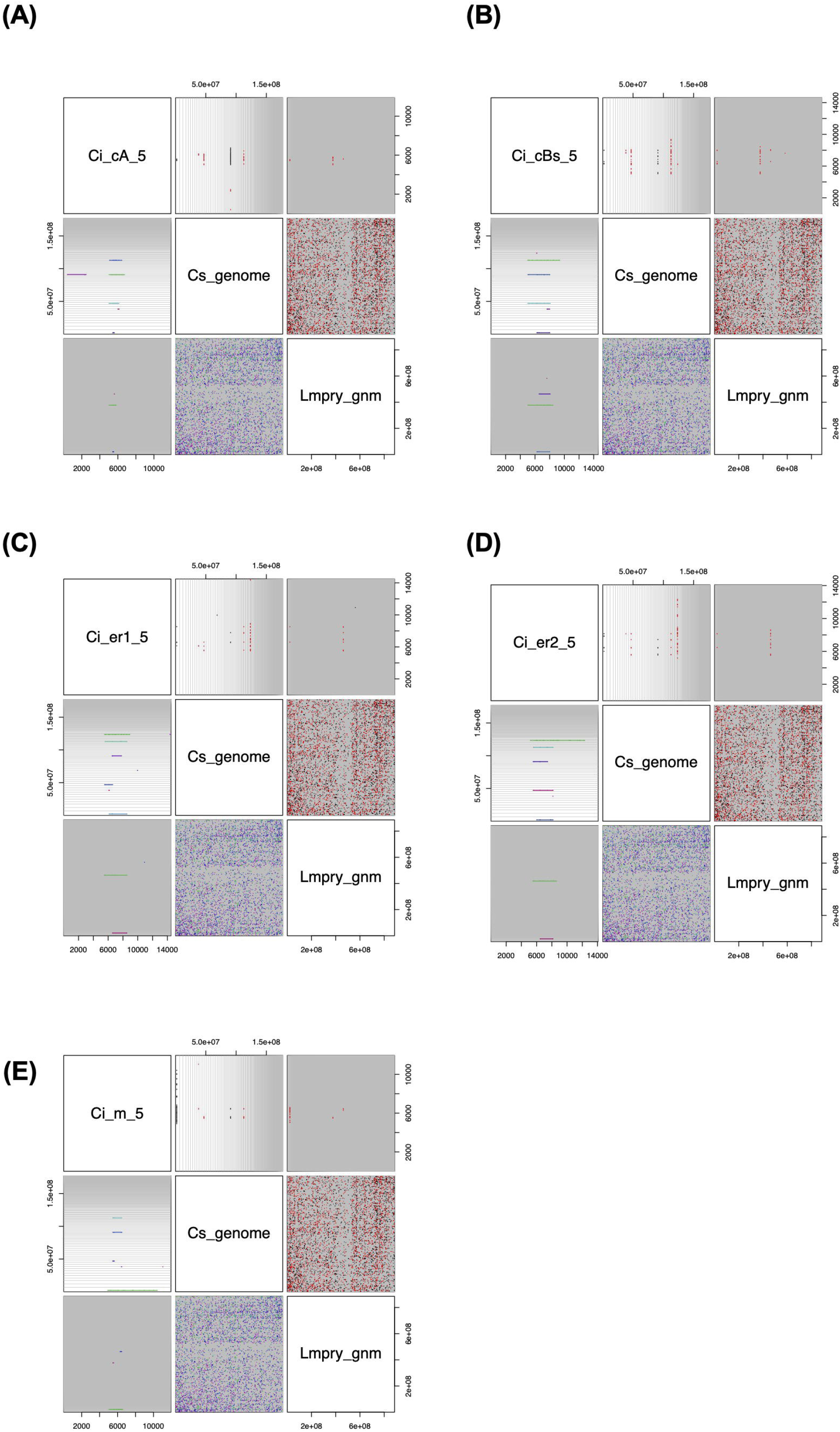
Synteny analysis between *C. intestinalis* genomic regions containing the HSP70 genes and the whole genome of *C. savignyi* and *P. marinus*. HSP70 gene regions were mapped to these genomes with 5 kb upstream and downstream flanking regions. The R DECIPHER package was used with default settings. Cs genome = Ciona_savignyi.CSAV2.0.dna.toplevel.fa. Pm genome = Petromyzon_marinus.Pmarinus_7.0.dna.toplevel.fa. (A) HSP70cA1. (B) HSP70cB1. (C) HSP70er1. (D) HSP70er2. (E) HSP70m1.

**Fig. 8.**
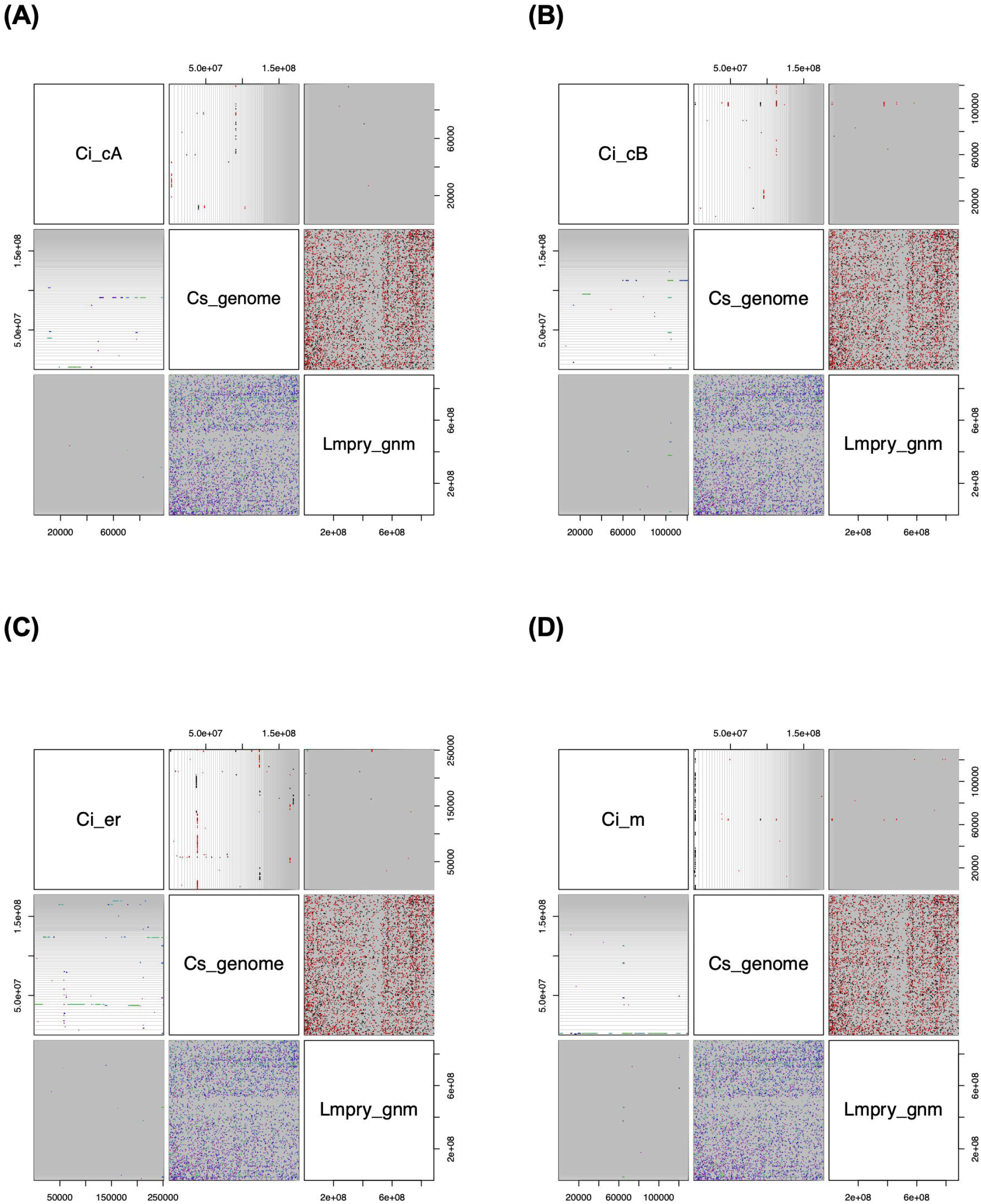
Synteny analysis between *C. intestinalis* genomic regions containing the HSP70 genes and the whole genome of *C. savignyi* and *P. marinus*. Regions around the HSP70 genes in Fig. 6 were mapped to these genomes by the DECIPHER package with default settings. Cs genome = Ciona_savignyi.CSAV2.0.dna.toplevel.fa. Pm genome = Petromyzon_marinus.Pmarinus_7.0.dna.toplevel.fa. (A) HSP70cA1, *C. intestinalis* KH:HT000129.1:1:98070:1. (B) HSP70cB1, *C. intestinalis* KH:HT000062.1:422312:543332:1. (C) HSP70er1, *C. intestinalis* KH:9:4803533:5054713:1. (D) HSP70m1, Ci Chr. 3 = *C. intestinalis* KH:3:5696489:5825960:1.

The results of synteny analysis suggest that there was a dynamic chromosomal rearrangement in the vertebrate lineage after the split from the common ancestor with tunicates, which makes it difficult to accurately trace the molecular evolutionary history of HSP70 genes in early vertebrates. However, it is likely that the regions containing the HSP70cA1 gene was the subject of extensive chromosomal rearrangement compared to those containing other HSP70 genes, which may explain the newly identified evolutionary event, the loss of HSP70cA gene in the vertebrate lineage.

## 4. Conclusion

In the present study, we demonstrated the presence of the HSP70cA genes in Chordata but did not find any evidence for their presence in Craniata. The synteny analysis suggests the loss of HSP70cA gene in early vertebrates, possibly caused by the chromosomal rearrangement. Despite the critical importance, the loss of HSP70 gene may not be a rare event in molecular evolution as suggested by the loss of HSP70cA, presence of HSP70m pseudogene in *C. savignyi*, and reported loss of HSP70 gene in hyperthermophilic archaea.

