## Supplementary Table 1 for "Genomic loss of the HSP70cA gene in the vertebrate lineage"

Supplementary Table S1. Keywords used for the NCBI database search

| Keyword | All filters | Number of hits |
|---|---|---|
| HSP70 | "HSP70"[All Fields] AND (animals[filter] AND (biomol_genomic[PROP] OR biomol_mrna[PROP]) AND ("1500"[SLEN] : "2500"[SLEN])) | 3,345 |
| HSC70 | "HSC70"[All Fields] AND (animals[filter] AND (biomol_genomic[PROP] OR biomol_mrna[PROP]) AND ("1500"[SLEN] : "2500"[SLEN])) | 290 |
| HSC71 | "HSC71"[All Fields] AND (animals[filter] AND (biomol_genomic[PROP] OR biomol_mrna[PROP]) AND ("1500"[SLEN] : "2500"[SLEN])) | 29 |
| "heat shock protein" 70 | "heat shock protein"[All Fields] AND 70[All Fields] AND (animals[filter] AND (biomol_genomic[PROP] OR biomol_mrna[PROP]) AND ("1500"[SLEN] : "2500"[SLEN])) | 1,051 |
| "heat shock cognate" | "heat shock cognate"[All Fields] AND (animals[filter] AND (biomol_genomic[PROP] OR biomol_mrna[PROP]) AND ("1500"[SLEN] : "2500"[SLEN])) | 883 |
| "heat shock cognate" 70 | "heat shock cognate"[All Fields] AND 70[All Fields] AND (animals[filter] AND (biomol_genomic[PROP] OR biomol_mrna[PROP]) AND ("1500"[SLEN] : "2500"[SLEN])) | 296 |
| "heat shock cognate" 71 | "heat shock cognate"[All Fields] AND 71[All Fields] AND (animals[filter] AND (biomol_genomic[PROP] OR biomol_mrna[PROP]) AND ("1500"[SLEN] : "2500"[SLEN])) | 552 |
| "stress 70" | "stress 70"[All Fields] AND (animals[filter] AND (biomol_genomic[PROP] OR biomol_mrna[PROP]) AND ("1500"[SLEN] : "2500"[SLEN])) | 126 |
