## Supplementary Table 2 for "Genomic loss of the HSP70cA gene in the vertebrate lineage"

airwise comparison

| Scientific name | Protein name (synonym) | Protein ID* |
|---|---|---|
| *N. vectensis* | HSP70cA1 (HSC71-1) | XP_001636593.1 |
| *A. millepora* | HSP70cA1 (HSP68) | XP_029210797.1 |
| *Priapulus* | HSP70cA1 (HSP70 A1-like) | XP_014681862.1 |
| *D. melanogaster* | HSP70cA1i (HSP70Aa) | AAG26887.1 |
| *D. melanogaster* | HSP70cA2i (HSP70Bb) | AAG26911.1 |
| *Bombyx* | HSP70cA1i | BAF69068.1 |
| *Spodoptera* | HSP70cA1i | ACN78407.1 |
| *Lingula* | HSP70cA1 (HSP70 1B-like) | XP_013382454.1 |
| *A. vaga* | HSP70cA2 (HSP70-1B) | GSADVT00016830001 (Genoscope) |
| *A. vaga* | HSP70cA1 (HSP70-1A) | GSADVT00001811001 (Genoscope) |
| *B. ibericus* | HSP70cA1i | ADR79281.1 |
| *Paralvinella* | HSP70cA1i (HSP70 form 1) | ABU63808.1 |
| *Aplysia* | HSP70cA1 (HSP70 B2-1) | XP_005103834.1 |
| *Aplysia* | HSP70cA2 (HSP70 B2-2) | XP_005100354.1 |
| *Crassostrea* HSP70 B2 | HSP70cA1 | EKC30019.1 |
| *Lottia* HSP70-1 (hypothetical protein LOTGIDRAFT_181897) | HSP70cA1 | XP_009051716.1 |
| *Lottia* HSP70-3 (hypothetical protein LOTGIDRAFT_209056) | HSP70cA3 | XP_009051717.1 |
| *Lottia* HSP70-2 (hypothetical protein LOTGIDRAFT_198956) | HSP70cA2 | XP_009045593.1 |
| *Lottia* HSP70-4 (hypothetical protein LOTGIDRAFT_190284) | HSP70cA4 | XP_009056460.1 |
| *Haliotis* heat shock inducible protein 70 | HSP70cA1 | ACO36048.1 |
| *A. queenslandica* HSC71-1 | HSP70cB1 | XP_011404208.1 |
| *A. queenslandica* HSC71-2 | HSP70cB2 | XP_011404198.1 |
| *A. millepora* HSC71-1 | HSP70cB1 | XP_029206535.1 |
| *A. millepora* HSC71-2 | HSP70cB2 | XP_029194855.1 |
| *N. vectensis* HSC71-2 | HSP70cB1 | XP_001622423.1 |
| *N. vectensis* HSC71-3 | HSP70cB2 | XP_001629343.1 |
| *H. vulgaris* HSP70 | HSP70cB1 | NP_001296624.1 |
| *Priapulus* HSPc | HSP70cB1 | XP_014668099.1 |
| *Priapulus* HSP70 II-like | HSP70cB2 | XP_014673112.1 |
| *Trichuris* HSP70 | HSP70cB1 | KFD56098.1 |
| *Daphnia* HSP70Bbb-1 | HSP70cB1 | KZS09073.1 |
| *Daphnia* HSP70Bbb-2 | HSP70cB2 | KZS10545.1 |
| *M. nipponense* HSP70-1 | HSP70cB1i | AGM50430.1 |
| *M. nipponense* HSC70 | HSP70cB2i | ABG45886.1 |
| *Limunus* HSP cognate 4 | HSP70cB1 | XP_013778668.1 |
| *Lingula* HSP71c | HSP70cB1 | XP_013380918.1 |

| | | |
|---|---|---|
| *Tevnia* HSP70-1 | HSP70cB1 | CBM42052.1 |
| *Tevnia* HSP70-2 | HSP70cB2 | CBM42053.1 |
| *Perinereis* HSP70-1 | HSP70cB1i | AND99892.1 |
| *Perinereis* HSP70-2 | HSP70cB2 | ADR66514.1 |
| *Paralvinella* HSP70 form 2 | HSP70cB1i | ABU63809.1 |
| *Helobdella* hypothetical protein 2 HELRODRAFT | HSP70cB1 | XP_009022152.1 |
| *Helobdella* hypothetical protein HELRODRAFT | HSP70cB2 | XP_009029445.1 |
| *Lottia* HSC70 (hypothetical protein LOTGIDRAFT_177837) | HSP70cB1 | XP_009046364.1 |
| *Aplysia* HSC71 | HSP70cB1 | XP_005098012.1 |
| *Haliotis* HSC70 | HSP70cB1 | ACO36047.1 |
| *Crassostrea* HSP71c | HSP70cB1 | BAD15287.1 |
| *Mytilus* HSP70 | HSP70cB1i | AAW52766.1 |
| *Mytilus* HSP71c | HSP70cB2 | CAH04109.1 |
| *Octopus* HSC71 | HSP70cB1 | XP_014783074.1 |
| *Octopus* HSP71-like | HSP70cB2 | XP_014784774.1 |
| *Oncorhynchus* HSC71 | HSP70cB1c | AAB21658.1 |
| *Alligator* HSP70A | HSP70cB1 | BAF94142.1 |
| *Pelodiscus* HSC70 | HSP70cB1i | ADO17794.1 |
| *Bos* HSP70 | HSP70cB2i | AAA73914.1 |
| *H. sapiens* HSP70-1B (HSPA1B) | HSP70cB2i | NP_005337.2 |
| *Rattus* HSP70 1B | HSP70cB2i | NP_114177.2 |
| *H. sapiens* HSP70-like1 (HSPA1L) | HSP70cB3c | NP_005518.3 |
| *H. sapiens* HSP70-6 (HSPA6) | HSP70cB5i | NP_002146.2 |
| *X. maculatus* HSP70-1 | HSP70cB1i | BAB72167.1 |
| *X. maculatus* HSP70-2 | HSP70cB2i | BAB72168.1 |
| *H. sapiens* HSP70-like2 (HSPA2) | HSP70cB4c | NP_068814.2 |
| *X. maculatus* HSC70 | HSP70cB3c | BAB72169.1 |
| *H. sapiens* HSC71 (HSPA8) | HSP70cB6c | NP_006588.1 |
| *Bos* HSC71 | HSP70cB1 | P19120.2 |
| *Rattus* HSC71 | HSP70cB1c | NP_077327.1 |

accession numbers are available in Yu et al. (2021)
