## Supplementary Table 3 for "Genomic loss of the HSP70cA gene in the vertebrate lineage"

Supplementary Table S3. Manual annotation of the 597 HSP70 genes in clusters 3 and 4 of Fig. 1C

| Cluster | Annotation[1] | DNA accession | Protein accession | Scientific name | Phylum | Class | Ser | Sequences around HSP70cA-specific Ser | Note |
|---|---|---|---|---|---|---|---|---|---|
| 3 | | Y15109.1 | CAA75383.1 | *Sycon raphanus* | Porifera | Calcarea | No | DD-GIFGVKSTAGD | |
| 3 | | X94985.1 | CAA64441.1 | *Geodia cydonium* | Porifera | Demospongiae | No | EE-GIFEVKSTAGD | |
| 3 | | XM_002114250.1 | XP_002114286.1 | *Trichoplax adhaerens* | Placozoa | - | No | D-NGIFEVKATAGD | No class found; subclass Testudinata |
| 3 | | HM147130.1 | ADM13667.1 | *Seriatopora hystrix* | Cnidaria | Anthozoa | No | DD-GIFEVKSTHGD | |
| 3 | HSC70cB1 | XM_001622373.2 | XP_001622423.1 | *Nematostella vectensis* | Cnidaria | Anthozoa | No | -EDGIFEVKSTHGD | |
| 3 | HSC70cB2 | XM_001629293.2 | XP_001629343.1 | *Nematostella vectensis* | Cnidaria | Anthozoa | No | -EDGIFEVKSTAGD | |
| 3 | | XM_015898489.1 | XP_015753975.1 | *Acropora digitifera* | Cnidaria | Anthozoa | No | -DGGIFEVKATRGV | |
| 3 | | XM_015898490.1 | XP_015753976.1 | *Acropora digitifera* | Cnidaria | Anthozoa | No | -DGGIFEVKATRGD | |
| 3 | | XM_015898491.1 | XP_015753977.1 | *Acropora digitifera* | Cnidaria | Anthozoa | No | DDG-IFEVKATRGY | |
| 3 | | XM_015898495.1 | XP_015753981.1 | *Acropora digitifera* | Cnidaria | Anthozoa | No | DD-GIFEVKATRGD | |
| 3 | | XM_020745144.1 | XP_020600803.1 | *Orbicella Faveolata* | Cnidaria | Anthozoa | No | DD-GIFEVKSTHGD | |
| 3 | | XM_021061767.2 | XP_020917426.1 | *Exaiptasia Diaphana* | Cnidaria | Anthozoa | No | -EDGIFEVKSTSGN | |
| 3 | | XM_022938441.1 | XP_022794176.1 | *Stylophora pistillata* | Cnidaria | Anthozoa | No | DD-GIFEVKSTHGD | |
| 3 | | XM_027187700.1 | XP_027043501.1 | *Pocillopora damicornis* | Cnidaria | Anthozoa | No | -EDGIFEVKSTAGN | |
| 3 | | XM_027196790.1 | XP_027052591.1 | *Pocillopora damicornis* | Cnidaria | Anthozoa | No | D-GIFEVKSTCGD | |
| 3 | | XM_028537258.1 | XP_028393059.1 | *Dendronephthya gigantea* | Cnidaria | Anthozoa | No | -EDGIFEVKSTAGD | |
| 3 | | XM_029339007.2 | XP_029194840.2 | *Acropora millepora* | Cnidaria | Anthozoa | No | -DGGIFEVKATRGD | |
| 3 | HSP70cB2 | XM_029339022.1 | XP_029194855.1 | *Acropora millepora* | Cnidaria | Anthozoa | No | DD-GIFEVKATRGD | |
| 3 | | XM_029339024.2 | XP_029194857.1 | *Acropora millepora* | Cnidaria | Anthozoa | No | DD-GIFEVKATRGD | |
| 3 | HSP70cB1 | XM_029350702.2 | XP_029206535.1 | *Acropora millepora* | Cnidaria | Anthozoa | No | -EDGIFEVKSTAGD | |
| 3 | | XM_044328936.1 | XP_044184871.1 | *Acropora millepora* | Cnidaria | Anthozoa | No | DD-GIFEVKATRGD | |
| 3 | | XM_044328958.1 | XP_044184893.1 | *Acropora millepora* | Cnidaria | Anthozoa | No | -DGGIFEVRATRGV | |
| 3 | | XM_044328963.1 | XP_044184898.1 | *Acropora millepora* | Cnidaria | Anthozoa | No | DDG-IFEVKATRGV | |
| 3 | | JN400918.1 | AEM62766.1 | *Hydractinia echinata* | Cnidaria | Hydrozoa | No | -EDGIFEVKSTHGD | |
| 3 | HSP70cB1 | NM_001309695.1 | NP_001296624.1 | *Hydra vulgaris* | Cnidaria | Hydrozoa | No | -EDGIFEVKSTAGD | |
| 3 | | Y09746.1 | CAA70893.1 | *Hydra oligactis* | Cnidaria | Hydrozoa | No | -EDGIFEVKSTAGD | |
| 3 | | U26448.1 | AAA99139.1 | *Echinococcus granulosus* | Platyhelminthes | Cestoda | No | -EDGIFEVKSTAGD | |
| 3 | | U70213.1 | AAB18390.1 | *Mesocestoides corti* | Platyhelminthes | Cestoda | No | -EDGIFEVKSTAGD | |
| 3 | | XM_024499810.1 | XP_024345771.1 | *Echinococcus granulosus* | Platyhelminthes | Cestoda | No | -EDGIFEVKSTAGD | |
| 3 | | KP027180.1 | AJN00640.1 | *Polycelis sp.* | Platyhelminthes | Rhabditophora | No | -EDGIFEVKSTAGD | |
| 3 | | KT899328.1 | ALJ32180.1 | *Schistosoma japonicum* | Platyhelminthes | Trematoda | No | -EDGIFEVKSTAGD | |
| 3 | | X05384.1 | CAA28976.1 | *Schistosoma mansoni* | Platyhelminthes | Trematoda | No | ED-GIFEVKSTAGD | |
| 3 | | XM_012947058.2 | XP_012802512.2 | *Schistosoma haematobium* | Platyhelminthes | Trematoda | No | EDG-IFEVKSTAGD | |
| 3 | | EF552221.1 | ABV46675.1 | *Angiostrongylus vasorum* | Nematoda | Chromadorea | No | -EDGIFEVKSTAGD | |
| 3 | | Y13114.1 | CAA73574.1 | *Trichinella britovi* | Nematoda | Enoplea | No | -EDGIFEVKSTAGD | |
| 3 | HSP70cB2 | XM_014817626.1 | XP_014673112.1 | *Priapulus caudatus* | Priapulida | Priapulimorpha | No | E-EGIFEVKSTAGD | |
| 3 | | MF542317.1 | AXZ96464.1 | *Neoseiulus barkeri* | Arthopoda | Arachnida | No | -EDGVFEVKSTAGD | |
| 3 | | MN556969.1 | QKY59635.1 | *Neoseiulus barkeri* | Arthopoda | Arachnida | No | -EDGVFEVKSTAGD | |
| 3 | | KX695141.1 | APH81352.1 | *Tigriopus kingsejongensis* | Arthopoda | Hexanauplia | No | -EDGIFEVKSTAGD | |
| 3 | | KY807149.1 | ASB34118.1 | *Eurytemora pacifica* | Arthopoda | Hexanauplia | No | EE-GIFEVKSTAGD | |
| 3 | | KU613183.1 | ANQ44717.1 | *Galathea squamifera* | Arthopoda | Malacostraca | No | -EDGIFEVKSTAGD | |
| 3 | | MH846234.1 | QCC72758.1 | *Macrobrachium rosenbergii* | Arthopoda | Malacostraca | No | DD-GVFEVKSTAGD | |
| 3 | | GQ495084.1 | ADM83424.1 | *Panonychus citri* | Arthropoda | Arachnida | No | DD-GIFEVKSTAGD | |
| 3 | | KC335213.1 | AGQ50609.1 | *Neoseiulus cucumeris* | Arthropoda | Arachnida | No | -EDGVFEVKSTAGD | |
| 3 | | MN556968.1 | QKY59634.1 | *Neoseiulus barkeri* | Arthropoda | Arachnida | No | DEG-VFEVKSTAGN | |
| 3 | | XM_027338446.1 | XP_027194247.1 | *Dermatophagoides pteronyssinus* | Arthropoda | Arachnida | No | DD-GIFEVKSTCGD | |
| 3 | | AF427596.1 | AAL27404.1 | *Artemia franciscana* | Arthropoda | Branchiopoda | No | -EDGIFEVKSTAGD | |
| 3 | | JX624124.1 | AGT28484.1 | *Pseudodiaptomus annandalei* | Arthropoda | Hexanauplia | No | EE-GIFEVKSTAGD | |
| 3 | | KF516629.1 | AII16533.1 | *Paracyclopina nana* | Arthropoda | Hexanauplia | No | -EDGIFEVKSTAGD | |
| 3 | | LT844577.1 | SMN34117.1 | *Lepeophtheirus salmonis* | Arthropoda | Hexanauplia | No | DD-GVFEVKSTAGD | |
| 3 | | LT844578.1 | SMN34118.1 | *Lepeophtheirus salmonis* | Arthropoda | Hexanauplia | No | -EDGVFEVKSTAGD | |
| 3 | | XM_040720011.1 | XP_040575945.1 | *Lepeophtheirus salmonis* | Arthropoda | Hexanauplia | No | DD-GVFEVKSTAGD | |
| 3 | | XM_040723696.1 | XP_040579630.1 | *Lepeophtheirus salmonis* | Arthropoda | Hexanauplia | No | -EDGIFEVKSTAGD | |
| 3 | | AH001015.2 | AAA28632.1 | *Drosophila simulans* | Arthropoda | Insecta | No | ED--GFEVKATAGD | |
| 3 | | FJ386398.1 | ACJ12782.1 | *Pyrrhocoris apterus* | Arthropoda | Insecta | - | – | Partial sequence without the Ser-containing region; Not used included in |
| 3 | | KC844150.1 | AGQ50301.1 | *Leguminivora glycinivorella* | Arthropoda | Insecta | No | -EDGIFEVKSTAGD | |
| 3 | | MG992397.1 | AXY94835.1 | *Galleria mellonella* | Arthropoda | Insecta | No | -EDGIFEVKSTAGD | |
| 3 | | MW456683.1 | UAJ82472.1 | *Pentalonia nigronervosa* | Arthropoda | Insecta | Yes | DEGSIFEVKSTAGD | Included in Fig. 3 |
| 3 | | NM_079339.3 | NP_524063.1 | *Drosophilia melanogaster* | Arthropoda | Insecta | No | -EDGIFEVKATAGD | |
| 3 | | XM_025551239.1 | XP_025407024.1 | *Sipha flava* | Arthropoda | Insecta | Yes | DEGSIFEVKSTAGD | Included in Fig. 3 |
| 3 | | XM_043393103.1 | XP_043249038.1 | *Colletes gigas* | Arthropoda | Insecta | No | N-DGIFEVKATAGD | |
| 3 | | AM410071.1 | CAL68987.1 | *Bythograea thermydron* | Arthropoda | Malacostraca | No | DD-GVFEVKSTAGD | |
| 3 | | AM410072.1 | CAL68988.1 | *Bythograea thermydron* | Arthropoda | Malacostraca | No | DDG-VFEGKSTAGD | |
| 3 | | AM410073.1 | CAL68989.1 | *Cyanagraea praedator* | Arthropoda | Malacostraca | No | DD-GVFEVKSTAGD | |
| 3 | | AM410074.1 | CAL68990.1 | *Segonzacia mesatlantica* | Arthropoda | Malacostraca | No | DD-GVFEVKSTAGD | |
| 3 | | AM410075.1 | CAL68991.1 | *Demania scaberrima* | Arthropoda | Malacostraca | No | DD-GVFEVKSTAGD | |
| 3 | | AM410076.1 | CAL68992.1 | *Xantho incisus* | Arthropoda | Malacostraca | No | DD-GVFEVKSTAGD | |
| 3 | | AM410079.1 | CAL68995.1 | *Cancer pagurus* | Arthropoda | Malacostraca | No | -EDGIFEVKSTAGD | |
| 3 | | DQ301506.1 | ABC01063.1 | *Procambarus clarki* | Arthropoda | Malacostraca | No | DEG-VFEVKSTAGD | |
| 3 | | DQ534065.1 | ABF85673.1 | *Rimicaris exoculata* | Arthropoda | Malacostraca | No | DD-GIFEVKSTAGD | |
| 3 | | FJ268954.1 | ACL52279.1 | *Rimicaris exoculata* | Arthropoda | Malacostraca | No | DD-GIFEVKSTAGD | |
| 3 | | FJ875280.1 | ACR54098.1 | *Palaemon varians* | Arthropoda | Malacostraca | No | DD-GVFEVKSTAGD | |
| 3 | | HG809965.1 | CDL93417.1 | *Gammarus pulex* | Arthropoda | Malacostraca | No | E-EGIFEVKSTAGD | |
| 3 | | HG809966.1 | CDL93418.1 | *Gammarus pulex* | Arthropoda | Malacostraca | No | DEG-IFEVKSTAGD | |
| 3 | | KC907348.1 | AGV07535.1 | *Chaceon affinis* | Arthropoda | Malacostraca | No | DEG-VFEVKSTAGD | |
| 3 | | KM067142.1 | AIR72268.1 | *Euphausia crystallorophias* | Arthropoda | Malacostraca | No | DD-GIFEVKATAGD | |
| 3 | | KM067147.1 | AIR72273.1 | *Euphausia superba* | Arthropoda | Malacostraca | No | DD-GIFEVKATAGD | |
| 3 | | KU613102.1 | ANQ44636.1 | *Maja squinado* | Arthropoda | Malacostraca | No | -EDGIFEVKSTAGD | |
| 3 | | KU613121.1 | ANQ44655.1 | *Homarus americanus* | Arthropoda | Malacostraca | No | DEG-VFEVKSTAGD | |
| 3 | | KU613122.1 | ANQ44656.1 | *Homarus americanus* | Arthropoda | Malacostraca | No | DEG-VFEVKSTAGD | |
| 3 | | KU613129.1 | ANQ44663.1 | *Astacus astacus* | Arthropoda | Malacostraca | No | DEG-VFEVKSTAGD | |
| 3 | | KU613139.1 | ANQ44673.1 | *Perisesarma bidens* | Arthropoda | Malacostraca | No | DD-GVFEVKATAGD | |
| 3 | | KU613143.1 | ANQ44677.1 | *Bythograea thermydron* | Arthropoda | Malacostraca | No | DD-GVFEVKSTAGD | |
| 3 | | KU613145.1 | ANQ44679.1 | *Segonzacia mesatlantica* | Arthropoda | Malacostraca | No | DD-GVFEVKSTAGD | |
| 3 | | KU613148.1 | ANQ44682.1 | *Homarus americanus* | Arthropoda | Malacostraca | No | DD-GIFEVKSTAGD | |
| 3 | | KU613149.1 | ANQ44683.1 | *Homarus gammarus* | Arthropoda | Malacostraca | No | DD-GIFEVKSTAGD | |
| 3 | | KU613150.1 | ANQ44684.1 | *Nephrops norvegicus* | Arthropoda | Malacostraca | No | DD-GIFEVKSTAGD | |
| 3 | | KU613151.1 | ANQ44685.1 | *Enoplometopus debelius* | Arthropoda | Malacostraca | No | DD-GIFEVKSTAGD | |
| 3 | | KU613153.1 | ANQ44687.1 | *Galathea strigosa* | Arthropoda | Malacostraca | No | DD-GIFEVKSTAGD | |
| 3 | | KU613154.1 | ANQ44688.1 | *Galathea squamifera* | Arthropoda | Malacostraca | No | DD-GIFEVKSTAGD | |
| 3 | | KU613169.1 | ANQ44703.1 | *Bythograea thermydron* | Arthropoda | Malacostraca | No | DEG-VFEVKSTAGD | |
| 3 | | KU613172.1 | ANQ44706.1 | *Perisesarma bidens* | Arthropoda | Malacostraca | No | DEG-VFEVKSTAGD | |
| 3 | | KU613174.1 | ANQ44708.1 | *Pachygrapsus marmoratus* | Arthropoda | Malacostraca | No | DEG-VFEVKSTAGD | |
| 3 | | KU613175.1 | ANQ44709.1 | *Hemigrapsus nudus* | Arthropoda | Malacostraca | No | DEG-VFEVKSTAGD | |
| 3 | | KU613176.1 | ANQ44710.1 | *Cardisoma armatum* | Arthropoda | Malacostraca | No | DEG-VFEVKSTAGD | |
| 3 | | KU613177.1 | ANQ44711.1 | *Hemigrapsus sanguineus* | Arthropoda | Malacostraca | No | DEG-VFEVKSTAGD | |
| 3 | | KU613187.1 | ANQ44721.1 | *Galathea squamifera* | Arthropoda | Malacostraca | No | DD-GIFEVKSTAGD | |
| 3 | | KU613190.1 | ANQ44724.1 | *Cancer pagurus* | Arthropoda | Malacostraca | No | DEG-VFEVKSTAGD | |
| 3 | | KU613191.1 | ANQ44725.1 | *Cancer pagurus* | Arthropoda | Malacostraca | No | DEG-VFEVKSTAGD | |
| 3 | | KX450403.1 | AQL57187.1 | *Thysanoessa inermis* | Arthropoda | Malacostraca | No | DD-GVFEVKATAGD | |
| 3 | | KX450404.1 | AQL57188.1 | *Thysanoessa inermis* | Arthropoda | Malacostraca | No | -EDGIFEVKSTAGD | |
| 3 | | MK750015.1 | QEJ88143.1 | *Macrophthalmus japonicus* | Arthropoda | Malacostraca | No | DD-GVFEVKSTAGD | |
| 3 | | XM_018151508.1 | XP_018006997.1 | *Hyalella azteca* | Arthropoda | Malacostraca | No | DD-GIFEVKATAGD | |
| 3 | HSP70cB1 | XM_009023904.1 | XP_009022152.1 | *Helobdella robusta* | Annelida | Clitellata | No | V-EGIFEVKSTSGD | |
| 3 | | XM_033768524.1 | XP_033624415.1 | *Asterias rubens* | Echinodermata | Asteroidea | No | -EDGIFEVKSTAGD | Included in Fig. 3 |
| 3 | | XM_033769388.1 | XP_033625279.1 | *Asterias rubens* | Echinodermata | Asteroidea | No | IEDGIFEVKSTAGD | Included in Fig. 3 |
| 3 | | AB062116.1 | BAB72170.1 | *Danio rerio* | Chordata | Actinopteri | No | -EDGIFEVKATAGD | |
| 3 | | MF770310.1 | AXC07422.1 | *Gadus chalcogrammus* | Chordata | Actinopteri | No | -EDGIFEVKSTAGD | |
| 3 | | NM_001104800.1 | NP_001098270.1 | *Oryzias latipes* | Chordata | Actinopteri | No | -EDGIFEVKSTAGD | |
| 3 | | XM_010789665.1 | XP_010787967.1 | *Notothenia coriiceps* | Chordata | Actinopteri | - | – | Predicted sequence with a calculated molecular weight 45890; possible gene prediction error; Not included in Fig. 2 |
| 3 | | XM_015337638.1 | XP_015193124.1 | *Lepisosteus oculatus* | Chordata | Actinopteri | - | – | Predicted sequence with a calculated molecular weight 43519; possible gene prediction error; Not included in Fig. 2 |

| | | | | | | | | | |
|---|---|---|---|---|---|---|---|---|---|
| 3 | | XM_019886392.1 | XP_019741951.1 | *Hippocampus comes* | Chordata | Actinopteri | No | DIDANGILNVSAVD | |
| 3 | | XM_022198378.1 | XP_022054070.1 | *Acanthochromis polyacanthus* | Chordata | Actinopteri | No | -EDGIFEVKSTAGD | |
| 3 | | XM_028579525.1 | XP_028435326.1 | *Perca flavescens* | Chordata | Actinopteri | No | -EDGIFEVKSTAGD | |
| 3 | | XM_033976677.1 | XP_033832568.1 | *Periophthalmus magnuspinnatus* | Chordata | Actinopteri | No | DEG-IFEVKATAGD | |
| 3 | | XM_034122700.1 | XP_033978591.1 | *Trematomus bernacchii* | Chordata | Actinopteri | No | IEDGIFEVKSTAGD | |
| 3 | | XM_038712511.1 | XP_038568439.1 | *Micropterus salmoides* | Chordata | Actinopteri | No | -EDGIFEVKSTAGD | |
| 3 | | XM_042481942.1 | XP_042337876.1 | *Plectropomus leopardus* | Chordata | Actinopteri | No | -EDGIFEVKSTAGD | |
| 3 | | XM_042707563.1 | XP_042563497.1 | *Clupea harengus* | Chordata | Actinopteri | No | -EDGIFEVKATAGD | |
| 3 | | MF405200.1 | ASS36960.1 | *Mugilogobius chulae* | Chordata | Actinopterygii | No | DD-GIFEVKATAGD | |
| 3 | | X71951.1 | CAA50749.1 | *Pleurodeles waltl* | Chordata | Amphibia | No | DD-GIFEVKATAGD | |
| 3 | | Y13661.1 | CAA74012.1 | *Pleurodeles waltl* | Chordata | Amphibia | No | -EDGIFEVKSTAGD | |
| 3 | | XM_039395228.1 | XP_039251162.1 | *Styela clava* | Chordata | Ascidiacea | No | DD-GIFEVKSTRGD | |
| 3 | | XM_039400825.1 | XP_039256759.1 | *Styela clava* | Chordata | Ascidiacea | No | DD-GIFEVKSTRGD | |
| 3 | | XM_039400826.1 | XP_039256760.1 | *Styela clava* | Chordata | Ascidiacea | No | DD-GIFEVKSTRGD | |
| 3 | | GU980869.1 | ADF81059.1 | *Gallus gallus* | Chordata | Aves | No | -EDGIFELKSTAGD | |
| 3 | | XM_007884395.2 | XP_007882586.2 | *Callorhinchus milii* | Chordata | Chondrichthyes | No | ENG-IFEVKSTAGD | |
| 3 | | HQ213893.1 | AEM65178.1 | *Kryptolebias marmoratus* | Chordata | Craniata | No | MEDGVFEVKATAGD | |
| 3 | | JN616381.1 | AET85555.1 | *Capra hircus* | Chordata | Craniata | No | DA-GVFEVTATAGD | |
| 3 | | KC172645.1 | AGF90789.1 | *Sebastes schlegelii* | Chordata | Craniata | No | -EDGIFEVKATAGD | |
| 3 | | XM_033027512.1 | XP_032883403.1 | *Phocoena sinus* | Chordata | Elasmobranchii | No | -EDGIFEVKSTSGD | |
| 3 | | XM_033047407.1 | XP_032903298.1 | *Amblyraja radiata* | Chordata | Elasmobranchii | No | -EDGIFEVKSTTGD | |
| 3 | | XM_033047408.1 | XP_032903299.1 | *Amblyraja radiata* | Chordata | Elasmobranchii | No | DD-GIFEVKSTTGD | |
| 3 | | XM_033047409.1 | XP_032903300.1 | *Amblyraja radiata* | Chordata | Elasmobranchii | No | -EDGIFEVKSTTGD | |
| 3 | | XM_033047410.1 | XP_032903301.1 | *Amblyraja radiata* | Chordata | Elasmobranchii | No | DD-GIFEVKSTTGD | |
| 3 | | XM_042438942.1 | XP_042294876.1 | *Sceloporus undulatus* | Chordata | Lepidosauria | No | -EDGIFEVKSTAGD | |
| 3 | | XM_039504114.1 | XP_039360048.1 | *Mauremys reevesii* | Chordata | Testudinata | No | -EDGIFEVKSTAGD | No class found; subclass Testudinata |
| 3 | | XM_039504115.1 | XP_039360049.1 | *Mauremys reevesii* | Chordata | Testudinata | No | -EDGLFEVKSTAGN | No class found; subclass Testudinata |
| 3 | | XM_039504119.1 | XP_039360053.1 | *Mauremys reevesii* | Chordata | Testudinata | No | -EDGLFEVKSTAGN | No class found; subclass Testudinata |
| 3 | | XM_039518346.1 | XP_039374280.1 | *Mauremys reevesii* | Chordata | Testudinata | No | -EDGLFEVKSTAGN | No class found; subclass Testudinata |
| 3 | | XM_043502053.1 | XP_043357988.1 | *Dermochelys coriacea* | Chordata | Testudinata | No | -EDGIFEVKSTAGD | No class found; subclass Testudinata |
| 3 | | LZPO01104799.1 | OBS60781.1 | *Neotoma lepida* | Chordata | Mammalia | - | – | Partial sequence without the Ser-containing region; Not used included in |
| 3 | | M12571.1 | AAA57234.1 | *Mus musculus* | Chordata | Mammalia | No | DANGILNVTATDKS | |
| 3 | | NM_001314206.1 | NP_001301135.1 | *Capra hircus* | Chordata | Mammalia | No | DD-GIFEVKATAGD | |
| 3 | HSP70cB5 | NM_002155.5 | NP_002146.2 | *Homo sapiens* | Chordata | Mammalia | No | DA-GVFEVKATAGD | HSP70-6 |
| 3 | | X51757.1 | CAA36061.1 | *Homo sapiens* | Chordata | Mammalia | No | DA-GVFEVKATAGD | |
| 3 | | XM_001488139.6 | XP_001488189.2 | *Equus caballus* | Chordata | Mammalia | No | DA-GVFEVKATAGD | |
| 3 | | XM_002809884.4 | XP_002809930.1 | *Pongo abelii* | Chordata | Mammalia | No | DAG-VFEVKATAGD | |
| 3 | | XM_004479707.2 | XP_004479764.1 | *Dasypus novemcinctus* | Chordata | Mammalia | No | DAG-VFEVKATAGD | |
| 3 | | XM_004846954.3 | XP_004847011.1 | *Heterocephalus glaber* | Chordata | Mammalia | No | DDG-IFEVKATAGD | |
| 3 | | XM_006041134.2 | XP_006041196.2 | *Bubalus bubalis* | Chordata | Mammalia | No | DAG-VFEVKATAGD | |
| 3 | | XM_006173149.3 | XP_006173211.1 | *Camelus ferus* | Chordata | Mammalia | No | DAG-VFEVKATAGD | |
| 3 | | XM_006219215.3 | XP_006219277.1 | *Vicugna pacos* | Chordata | Mammalia | No | DA-GVFEVKATAGD | |
| 3 | | XM_007990048.2 | XP_007988239.2 | *Chlorocebus sabaeus* | Chordata | Mammalia | Yes | DGISEVKSTARARN | Included in Fig. 3 |
| 3 | | XM_008054287.1 | XP_008052478.1 | *Carlito syrichta* | Chordata | Mammalia | No | DA-GVFEVKATAGD | |
| 3 | | XM_011770692.2 | XP_011768994.2 | *Macaca nemestrina* | Chordata | Mammalia | No | EYG-IFEIKSTARD | |
| 3 | | XM_012001234.1 | XP_011856624.1 | *Mandrillus leucophaeus* | Chordata | Mammalia | No | EDG-ISEVKSTARG | There's a Ser but not an insersion |
| 3 | | XM_012449593.1 | XP_012305016.1 | *Aotus nancymaae* | Chordata | Mammalia | No | DA-GVFEVKATAGD | |
| 3 | | XM_012554558.2 | XP_012410012.2 | *Trichechus manatus latirostris* | Chordata | Mammalia | No | EDG-IFEVKSTAGY | |
| 3 | | XM_013004979.2 | XP_012860433.2 | *Echinops telfairi* | Chordata | Mammalia | No | EDG-IFEVKSTARD | |
| 3 | | XM_014584767.2 | XP_014440253.1 | *Tupaia chinensis* | Chordata | Mammalia | No | EDG-SFEVRSTAGD | There's a Ser but not an insersion |
| 3 | | XM_015133087.2 | XP_014988573.1 | *Macaca mulatta* | Chordata | Mammalia | No | -EYGIFEIKSTARD | |
| 3 | | XM_015446958.1 | XP_015302444.1 | *Macaca fascicularis* | Chordata | Mammalia | No | -EYGIFEIKSTARD | |
| 3 | | XM_015544778.1 | XP_015400264.1 | *Panthera tigris altaica* | Chordata | Mammalia | No | E-YGILEVKSTAGD | |
| 3 | | XM_015569997.1 | XP_015425483.1 | *Myotis davidii* | Chordata | Mammalia | No | VEDGIFEVKSTAGG | |
| 3 | | XM_017875616.1 | XP_017731105.1 | *Rhinopithecus bieti* | Chordata | Mammalia | No | GRDGIFEVNSTAGD | |
| 3 | | XM_020286087.1 | XP_020141676.1 | *Microcebus murinus* | Chordata | Mammalia | No | IENGIFEVKFTAGD | |
| 3 | | XM_020875037.1 | XP_020730696.1 | *Odocoileus virginianus texanus* | Chordata | Mammalia | No | DA-GVFEVKATAGD | |
| 3 | | XM_024128213.2 | XP_023983981.1 | *Physeter catodon* | Chordata | Mammalia | No | D-GISEVKSTAGD | |
| 3 | | XM_024129765.1 | XP_023985533.1 | *Physeter catodon* | Chordata | Mammalia | No | -EDGIFEVKSTAGD | |
| 3 | | XM_026381449.1 | XP_026237234.1 | *Urocitellus parryii* | Chordata | Mammalia | No | DA-GVFEVRATAGD | |
| 3 | | XM_027126839.1 | XP_026982640.1 | *Lagenorhynchus obliquidens* | Chordata | Mammalia | No | ED-GIFEVQSTDTH | |
| 3 | | XM_027126845.1 | XP_026982646.1 | *Lagenorhynchus obliquidens* | Chordata | Mammalia | No | ED-GIFEVQSTDTH | |
| 3 | | XM_027126854.1 | XP_026982655.1 | *Lagenorhynchus obliquidens* | Chordata | Mammalia | No | ED-GIFEVQSTDTH | |
| 3 | | XM_027977462.2 | XP_027833263.1 | *Ovis aries* | Chordata | Mammalia | No | DA-GVFEVKATAGD | |
| 3 | | XM_031435899.1 | XP_031291759.1 | *Camelus dromedarius* | Chordata | Mammalia | No | DA-GVFEVKATAGD | |
| 3 | | XM_032620774.1 | XP_032476665.1 | *Contarinia nasturtii* | Chordata | Mammalia | No | E-DGISEVKSTAGD | |
| 3 | | XM_032621674.1 | XP_032477565.1 | *Phocoena sinus* | Chordata | Mammalia | No | -EDGIFEVKSTAGD | |
| 3 | | XM_032637328.1 | XP_032493219.1 | *Phocoena sinus* | Chordata | Mammalia | No | -EDGIFEIQSTDTH | |
| 3 | | XM_033403021.1 | XP_033258912.1 | *Belonocnema treatae* | Chordata | Mammalia | No | -EDGIFEVKSTAGD | |
| 3 | | XM_033406920.1 | XP_033262811.1 | *Orcinus orca* | Chordata | Mammalia | No | -EDGIFEVKSTAGD | |
| 3 | | XM_033423668.1 | XP_033279559.1 | *Orcinus orca* | Chordata | Mammalia | No | ED-GIFEVQSTDTH | |
| 3 | | XM_033438257.1 | XP_033294148.1 | *Orcinus orca* | Chordata | Mammalia | No | -ENGIFEVKSTAGD | |
| 3 | | XM_033849067.1 | XP_033704958.1 | *Tursiops truncatus* | Chordata | Mammalia | No | E-DGISEVKSTAGD | |
| 3 | | XM_033849164.1 | XP_033705055.1 | *Tursiops truncatus* | Chordata | Mammalia | No | -EDGIFEVKSTAGD | |
| 3 | | XM_033852269.1 | XP_033708160.1 | *Tursiops truncatus* | Chordata | Mammalia | No | -EDGIFEVKSTAGD | |
| 3 | | XM_033859974.1 | XP_033715865.1 | *Tursiops truncatus* | Chordata | Mammalia | No | IEGGIFEVQSTDTH | |
| 3 | | XM_034498555.1 | XP_034354446.1 | *Arvicanthis niloticus* | Chordata | Mammalia | No | -EDGIFEVKSTAGD | |
| 3 | | XM_034963713.1 | XP_034819604.1 | *Pan paniscus* | Chordata | Mammalia | No | -EDGIFEVKSTAGD | |
| 3 | | XM_035727524.1 | XP_035583417.1 | *Zalophus californianus* | Chordata | Mammalia | No | -EDGIFEVKSTAG | |
| 3 | | XM_036091286.1 | XP_035947179.1 | *Halichoerus grypus* | Chordata | Mammalia | No | LEDGIFEVKSTAGD | |
| 3 | | XM_036454008.1 | XP_036309901.1 | *Pipistrellus kuhlii* | Chordata | Mammalia | No | -EDGIFEVKSTAGD | |
| 3 | | XM_036460795.1 | XP_036316688.1 | *Pipistrellus kuhlii* | Chordata | Mammalia | No | E-DGIFEVKSTAVD | |
| 3 | | XM_036844991.1 | XP_036700886.1 | *Balaenoptera musculus* | Chordata | Mammalia | No | IEGGIFEVKSTAGD | |
| 3 | | XM_036858029.1 | XP_036713924.1 | *Balaenoptera musculus* | Chordata | Mammalia | No | -EDGIFEVKSTDTH | |
| 3 | | XM_037157000.1 | XP_037012895.1 | *Artibeus jamaicensis* | Chordata | Mammalia | No | -EEGIFEVKATAGD | |
| 3 | | XM_037732726.1 | XP_037588654.1 | *Cebus imitator* | Chordata | Mammalia | No | -EDGIFEVKSTARD | |
| 3 | | XM_037813325.1 | XP_037669253.1 | *Choloepus didactylus* | Chordata | Mammalia | No | -DGGVFEVKATAGD | |
| 3 | | XM_037982818.1 | XP_037838746.1 | *Chlorocebus sabaeus* | Chordata | Mammalia | No | K-DGIFEIKSTARD | |
| 3 | | XM_038660500.1 | XP_038516428.1 | *Canis lupus familiaris* | Chordata | Mammalia | No | -EDGIFEVKSTAGD | |
| 3 | | XM_038770107.1 | XP_038626035.1 | *Tachyglossus aculeatus* | Chordata | Mammalia | No | -DGIFEVKSTAGD | |
| 3 | | XM_038770111.1 | XP_038626039.1 | *Tachyglossus aculeatus* | Chordata | Mammalia | No | -RYEILEVKSTAGD | |
| 3 | | XM_039475627.1 | XP_039331561.1 | *Saimiri boliviensis boliviensis* | Chordata | Mammalia | No | DD-GIFEVKSTRGD | |
| 3 | | XM_039479873.1 | XP_039335807.1 | *Saimiri boliviensis boliviensis* | Chordata | Mammalia | No | -EDGIFEVKSTAGD | |
| 3 | | XM_040446639.1 | XP_040302573.1 | *Puma yagouaroundi* | Chordata | Mammalia | No | TEDGIFEVKSTARD | |
| 3 | | XM_040475920.1 | XP_040331854.1 | *Puma yagouaroundi* | Chordata | Mammalia | No | -EDGIFEVKSTAGD | |
| 3 | | XM_040484222.1 | XP_040340156.1 | *Puma yagouaroundi* | Chordata | Mammalia | No | ED-GIFEVKSAGDT | |
| 3 | | XM_041740316.1 | XP_041596250.1 | *Vulpes lagopus* | Chordata | Mammalia | No | -EDGIFEVKSTAGD | |
| 3 | | XM_041763305.1 | XP_041619239.1 | *Vulpes lagopus* | Chordata | Mammalia | No | TEDGIFEVKSTAGD | |
| 3 | | XM_042237739.1 | XP_042093673.1 | *Ovis aries* | Chordata | Mammalia | No | -EDGIFEVKSTAGD | |
| 3 | | XM_042924886.1 | XP_042780820.1 | *Panthera leo* | Chordata | Mammalia | No | TEDGIFEVKSTAKD | |
| 3 | | XM_042975793.1 | XP_042831727.1 | *Panthera tigris* | Chordata | Mammalia | No | TEDGIFEVKSTAKD | |
| 3 | | XM_043568906.1 | XP_043424841.1 | *Prionailurus bengalensis* | Chordata | Mammalia | No | TEDGIFEVKSTARD | |
| 3 | | XM_044657370.1 | XP_044513305.1 | *Gracilinanus agilis* | Chordata | Mammalia | No | -EDGIFEVKSTAGD | |
| 4 | | HF570334.1 | CCQ18651.1 | *Sycon ciliatum* | Porifera | Calcerea | No | DN-GLFAVKSTAGD | Included in Fig. 3 |
| 4 | HSP70cB2 | XM_011405896.2 | XP_011404198.1 | *Amphimedon queenslandica* | Porifera | Demospongiae | No | -ENGIFEVKSIIGD | Included in Fig. 3 |
| 4 | HSP70cB1 | XM_011405906.2 | XP_011404208.1 | *Amphimedon queenslandica* | Porifera | Demospongiae | No | -EDGIFEVRSTSGN | Included in Fig. 3 |
| 4 | | Y13926.1 | CAA74243.1 | *Rhabdocalyptus dawsoni* | Porifera | Hexactinellida | No | -EDGVFEVRSTAGD | Included in Fig. 3 |
| 4 | | AB201749.1 | BAD89541.1 | *Pocillopora damicornis* | Cnidaria | Anthozoa | Yes | DEGSFFQVLSTAGN | |
| 4 | | KP330265.1 | AKC91104.1 | *Stylophora pistillata* | Cnidaria | Anthozoa | Yes | DEGSFFQVLSTAGN | |
| 4 | | KU902046.1 | AOF40009.1 | *Stylophora pistillata* | Cnidaria | Anthozoa | Yes | DEGSIFEVKATAGD | |
| 4 | | KU902047.1 | AOF40010.1 | *Stylophora pistillata* | Cnidaria | Anthozoa | Yes | DEGSIFEVKATAGD | |
| 4 | | KU920674.1 | AMX23290.1 | *Pocillopora damicornis* | Cnidaria | Anthozoa | Yes | DEGSIFEVKATAGD | |
| 4 | HSP70cA1 | XM_001636543.2 | XP_001636593.1 | *Nematostella vectensis* | Cnidaria | Anthozoa | Yes | DDGSLFEVKSTAGD | |
| 4 | | XM_020763579.1 | XP_020619238.1 | *Orbicella Faveolata* | Cnidaria | Anthozoa | Yes | DEGSFFEVQSTAGN | |
| 4 | | XM_022923313.1 | XP_022779048.1 | *Stylophora pistillata* | Cnidaria | Anthozoa | Yes | DEGSFFQVLSTAGN | |
| 4 | | XM_027181441.1 | XP_027037242.1 | *Pocillopora damicornis* | Cnidaria | Anthozoa | No | ED-GIFEVQSTDTH | |
| 4 | HSP70cA1 | XM_029354964.1 | XP_029210797.1 | *Acropora millepora* | Cnidaria | Anthozoa | Yes | DEGSLFEVRSTAGD | Included in Fig. 3 |
| 4 | | GL382985.1 | EGT52639.1 | *Caenorhabditis brenneri* | Nematoda | Chromadorea | Yes | DEGSLFEVRSTAGD | |

| | | | | | | | | | |
|---|---|---|---|---|---|---|---|---|---|
| 4 | | XM_003104036.1 | XP_003104084.1 | *Caenorhabditis remanei* | Nematoda | Chromadorea | Yes | SEGSIFEVKSTAGD | |
| 4 | | XM_003105102.1 | XP_003105150.1 | *Caenorhabditis remanei* | Nematoda | Chromadorea | Yes | AEGSIFEVRSTAGD | |
| 4 | HSP70cA1 | XM_014826376.1 | XP_014681862.1 | *Priapulus caudatus* | Priapulida | Priapulimorpha | No | DEGAMFEVRSTAGD | Ser is replaced by Ala as reported in Yu et al. (2021) |
| 4 | | AY651782.1 | AAT75324.1 | *Ixodes scapularis* | Arthropoda | Arachnida | Yes | DEGSMFEVRSTAGD | |
| 4 | | DQ778943.1 | ABG47443.1 | *Dermacentor variabilis* | Arthropoda | Arachnida | Yes | DEGSIFEVKATIAG | |
| 4 | | DQ778944.1 | ABG47444.1 | *Dermacentor variabilis* | Arthropoda | Arachnida | Yes | DEGSIFEVKATIAG | |
| 4 | | KY249381.1 | APO36715.1 | *Haemaphysalis flava* | Arthropoda | Arachnida | Yes | DQGSMFEVRSTAGD | |
| 4 | | MN518895.1 | QKR72386.1 | *Haemaphysalis flava* | Arthropoda | Arachnida | Yes | DEGSMFEVRSTAGD | |
| 4 | | MN820834.1 | QUQ60736.1 | *Pardosa pseudoannulata* | Arthropoda | Arachnida | Yes | DEGSLFEVRSTAGD | |
| 4 | | MT679220.1 | QNL15810.1 | *Dermacentor silvarum* | Arthropoda | Arachnida | Yes | DEGSMFEVRSTAGD | |
| 4 | | XM_016048359.3 | XP_015903845.1 | *Parasteatoda tepidariorum* | Arthropoda | Arachnida | Yes | DEGSLFEVRSTAGD | |
| 4 | | XM_016074530.3 | XP_015930016.2 | *Parasteatoda tepidariorum* | Arthropoda | Arachnida | Yes | DEGSLFEVRSTAGD | |
| 4 | | XM_035350231.1 | XP_035206122.1 | *Stegodyphus dumicola* | Arthropoda | Arachnida | Yes | DEGSLFEVKATSGD | |
| 4 | | XM_035378018.1 | XP_035233909.1 | *Stegodyphus dumicola* | Arthropoda | Arachnida | Yes | DEGSLFEVRSTAGD | |
| 4 | | XM_043363705.1 | XP_043219640.1 | *Amphibalanus amphitrite* | Arthropoda | Hexanauplia | Yes | ADGSMFEVLSTAGD | |
| 4 | | XM_043382268.1 | XP_043238203.1 | *Amphibalanus amphitrite* | Arthropoda | Hexanauplia | Yes | ADGSMFEVKSTAGD | |
| 4 | | XM_043386251.1 | XP_043242186.1 | *Amphibalanus amphitrite* | Arthropoda | Hexanauplia | Yes | ADGSMFEVKSTAGD | |
| 4 | HSP70cA1i | AB035326.1 | BAF69068.1 | *Bombyx mori* | Arthropoda | Insecta | Yes | DEGSLFEVKSTAGD | |
| 4 | | AB162946.1 | BAD42358.1 | *Chironomus yoshimatsui* | Arthropoda | Insecta | Yes | DEGSLFEVRSTAGD | |
| 4 | | AB179657.1 | BAD18974.1 | *Antheraea yamamai* | Arthropoda | Insecta | Yes | DEGSLFEVKSTAGD | |
| 4 | | AB251895.1 | BAF03555.1 | *Mamestra brassicae* | Arthropoda | Insecta | Yes | DEGSLFEVRATAGD | |
| 4 | | AB325801.1 | BAF95560.1 | *Plutella xylostella* | Arthropoda | Insecta | Yes | DEGSLFEVKSTAGD | |
| 4 | | AF107338.2 | AAD17995.2 | *Sarcophaga crassipalpis* | Arthropoda | Insecta | Yes | DEGSLFEVRATAGD | |
| 4 | | AF288978.2 | AAG01177.2 | *Leptinotarsa decemlineata* | Arthropoda | Insecta | Yes | DEGSLFEVKSTAGD | |
| 4 | HSP70cA1i | AF295933.1 | AAG26887.1 | *Drosophila melanogaster* | Arthropoda | Insecta | Yes | DEGSLFEVRSTAGD | |
| 4 | | AF295938.1 | AAG26892.1 | *Drosophila melanogaster* | Arthropoda | Insecta | Yes | DEGSLFEVRSTAGD | |
| 4 | | AF295942.1 | AAG26896.1 | *Drosophila melanogaster* | Arthropoda | Insecta | Yes | DEGSLFEVRSTAGD | |
| 4 | | AF295953.1 | AAG26907.1 | *Drosophila melanogaster* | Arthropoda | Insecta | Yes | DEGSLFEVRSTAGD | |
| 4 | | AF295955.1 | AAG26909.1 | *Drosophila melanogaster* | Arthropoda | Insecta | Yes | DEGSLFEVRSTAGD | |
| 4 | HSP70cA2i | AF295957.1 | AAG26911.1 | *Drosophila melanogaster* | Arthropoda | Insecta | Yes | DEGSLFEVRSTAGD | |
| 4 | | AF295958.1 | AAG26912.1 | *Drosophila melanogaster* | Arthropoda | Insecta | Yes | YEGSLFEVRSTAGD | |
| 4 | | AF295959.1 | AAG26913.1 | *Drosophila melanogaster* | Arthropoda | Insecta | Yes | DEGSLFEVRSTAGD | |
| 4 | | AF295962.1 | AAG26916.1 | *Drosophila melanogaster* | Arthropoda | Insecta | Yes | DEGSLFEVRSTAGD | |
| 4 | | AF295963.1 | AAG24834.1 | *Drosophila simulans* | Arthropoda | Insecta | Yes | DEGSLFEVRSTAGD | |
| 4 | | AF295964.1 | AAG24835.1 | *Drosophila simulans* | Arthropoda | Insecta | Yes | DEGSLFEVRSTAGD | |
| 4 | | AF295965.1 | AAG24836.1 | *Drosophila simulans* | Arthropoda | Insecta | Yes | DEGSLFEVRSTAGD | |
| 4 | | AF295966.1 | AAG24837.1 | *Drosophila simulans* | Arthropoda | Insecta | Yes | DEGSLFEVRSTAGD | |
| 4 | | AF295968.1 | AAG24839.1 | *Drosophila simulans* | Arthropoda | Insecta | Yes | DEGSLFEVRSTAGD | |
| 4 | | AF295969.1 | AAG24840.1 | *Drosophila simulans* | Arthropoda | Insecta | Yes | DEGSLFEVRSTAGD | |
| 4 | | AF295971.1 | AAG24842.1 | *Drosophila simulans* | Arthropoda | Insecta | Yes | DEGSLFEVRSTAGD | |
| 4 | | AF295972.1 | AAG24843.1 | *Drosophila simulans* | Arthropoda | Insecta | Yes | DEGSLFEVRSTAGD | |
| 4 | | AF295973.1 | AAG24844.1 | *Drosophila simulans* | Arthropoda | Insecta | Yes | DEGSLFEVRSTAGD | |
| 4 | | AF295974.1 | AAG24845.1 | *Drosophila simulans* | Arthropoda | Insecta | Yes | DEGSLFEVRSTAGD | |
| 4 | | AF295975.1 | AAG24846.1 | *Drosophila simulans* | Arthropoda | Insecta | Yes | DEGSLFEVRSTAGD | |
| 4 | | AF295976.1 | AAG24847.1 | *Drosophila simulans* | Arthropoda | Insecta | Yes | DEGSLFEVRSTAGD | |
| 4 | | AF295977.1 | AAG24848.1 | *Drosophila simulans* | Arthropoda | Insecta | Yes | DEGSLSEVRSTAGD | |
| 4 | | AF295978.1 | AAG24849.1 | *Drosophila simulans* | Arthropoda | Insecta | Yes | DEGSLFEVRSTAGD | |
| 4 | | AF302410.1 | AAG24874.1 | *Drosophila orena* | Arthropoda | Insecta | Yes | DEGSLFEVRSTAGD | |
| 4 | | AF302411.1 | AAG24875.1 | *Drosophila orena* | Arthropoda | Insecta | Yes | DEGSLFEVRSTAGD | |
| 4 | | AF302412.1 | AAG24876.1 | *Drosophila orena* | Arthropoda | Insecta | Yes | DEGSLFEVRSTAGD | |
| 4 | | AF302413.1 | AAG24877.1 | *Drosophila orena* | Arthropoda | Insecta | Yes | DEGSLFEVRSTAGD | |
| 4 | | AF302416.1 | AAG25967.1 | *Drosophila mauritiana* | Arthropoda | Insecta | Yes | DEGSLFEVRSTAGD | |
| 4 | | AF302420.1 | AAG25968.1 | *Drosophila mauritiana* | Arthropoda | Insecta | Yes | DEGSLFEVRSTAGD | |
| 4 | | AF302421.1 | AAG25969.1 | *Drosophila mauritiana* | Arthropoda | Insecta | Yes | DEGSLFEVRSTAGD | |
| 4 | | AF322911.1 | AAG42838.1 | *Leptinotarsa decemlineata* | Arthropoda | Insecta | Yes | DEGSLFEVRATAGD | |
| 4 | | AF350459.1 | AAK30216.1 | *Drosophila melanogaster* | Arthropoda | Insecta | Yes | DEGSLFEVRSTAGD | |
| 4 | | AF350485.1 | AAK30242.1 | *Drosophila melanogaster* | Arthropoda | Insecta | Yes | DEGSLFEVRSTAGD | |
| 4 | | AF350489.1 | AAK30246.1 | *Drosophila melanogaster* | Arthropoda | Insecta | Yes | DEGSLFEVRSTAGD | |
| 4 | | AF350490.1 | AAK30247.1 | *Drosophila melanogaster* | Arthropoda | Insecta | Yes | DEGSLFEVRSTAGD | |
| 4 | | AF350491.1 | AAK30248.1 | *Drosophila melanogaster* | Arthropoda | Insecta | Yes | DEGSLFEVRSTAGD | |
| 4 | | AJ001365.1 | CAA04699.1 | *Drosophila auraria* | Arthropoda | Insecta | Yes | DEGSLFEVRATAGD | |
| 4 | | AY163157.2 | AAN85117.1 | *Chironomus tentans* | Arthropoda | Insecta | Yes | DEGSLFEVRSTAGD | |
| 4 | | AY220911.1 | AAO65964.1 | *Manduca sexta* | Arthropoda | Insecta | Yes | DEGSLFEVKATAGD | |
| 4 | | AY842476.2 | AAW32098.2 | *Liriomyza huidobrensis* | Arthropoda | Insecta | Yes | DEGSLFEVRSTAGD | |
| 4 | | AY842477.2 | AAW32099.2 | *Liriomyza sativae* | Arthropoda | Insecta | Yes | DEGSLFEVRSTAGD | |
| 4 | | AY974355.1 | AAX84696.1 | *Culex pipiens* | Arthropoda | Insecta | Yes | DEGSLFEVRSTAGD | |
| 4 | | DQ017057.1 | AAY28732.1 | *Delia antiqua* | Arthropoda | Insecta | Yes | DEGSIFEVKATAGD | |
| 4 | | DQ093385.2 | ACZ52196.1 | *Bemisia tabaci* | Arthropoda | Insecta | Yes | DEGSIFEVKATAGD | |
| 4 | | EF103584.1 | ABL06948.1 | *Rhagoletis pomonella* | Arthropoda | Insecta | Yes | DEGSIFEVKATAGD | |
| 4 | | EF445946.1 | ABO31121.1 | *Lucilia cuprin* | Arthropoda | Insecta | Yes | DEGSLFEVRSTAGD | |
| 4 | | EF523381.1 | ABP93405.1 | *Omphisa fuscidentalis* | Arthropoda | Insecta | Yes | DEGSIFEVKATAGD | |
| 4 | | EF569673.1 | ABQ39970.1 | *Anatolica polita borealis* | Arthropoda | Insecta | Yes | DEGSIFEVKATAGD | |
| 4 | | EU137871.1 | ABV55505.1 | *Microplitis mediator* | Arthropoda | Insecta | Yes | DEGSIFEVKATAGD | |
| 4 | | EU430480.1 | ABZ10939.1 | *Sesamia nonagrioides botanephaga* | Arthropoda | Insecta | No | DEGALFEVRATAGD | |
| 4 | | EU523048.1 | ACB59073.1 | *Stratiomys singularior* | Arthropoda | Insecta | Yes | DEGSLFEVRATAGG | |
| 4 | | EU585779.1 | ACD84944.1 | *Macrocentrus cingulum* | Arthropoda | Insecta | Yes | DEGSLFEVKSAAGD | |
| 4 | | EU684308.1 | ACD63050.1 | *Exorista civilis* | Arthropoda | Insecta | Yes | DEGSIFEVKATAGD | |
| 4 | | EU861391.2 | ACF74975.2 | *Trialeurodes vaporariorum* | Arthropoda | Insecta | Yes | DEGSLFEVRSTAGD | |
| 4 | | EU934244.1 | ACH85201.1 | *Trialeurodes vaporariorum* | Arthropoda | Insecta | Yes | DEGSLFEVRSTAGD | |
| 4 | HSP70cA1i | FJ754276.1 | ACN78407.1 | *Spodoptera exigua* | Arthropoda | Insecta | Yes | DEGSLFEVRATAGD | |
| 4 | | FJ862049.1 | ACQ78180.1 | *Spodoptera exigua* | Arthropoda | Insecta | Yes | DEGSIFEVKATAGD | |
| 4 | | GAMD01002785.1 | JAA98805.1 | *Anopheles aquasalis* | Arthropoda | Insecta | Yes | DEGSLFEVRSTAGD | |
| 4 | | GQ199477.1 | ACS72236.1 | *Heliothis viriplaca* | Arthropoda | Insecta | Yes | DEGSLFEVRATAGD | |
| 4 | | GQ389711.1 | ACV32640.1 | *Helicoverpa zea* | Arthropoda | Insecta | Yes | DEGSLFEVRSTAGD | |
| 4 | | GU074513.1 | ACY71070.1 | *Bemisia tabaci* | Arthropoda | Insecta | Yes | DEGSLFEVRATAGD | |
| 4 | | GU591409.1 | ADQ12986.1 | *Bactrocera dorsalis* | Arthropoda | Insecta | Yes | DEGSLFGVRATAGD | |
| 4 | | GU945198.1 | ADI50267.1 | *Antheraea pernyi* | Arthropoda | Insecta | No | DEGSIFEVKATAGD | |
| 4 | | HM212645.1 | ADK39311.1 | *Plutella xylostella* | Arthropoda | Insecta | No | –EDGIFEVKSTAGD | |
| 4 | | HM370509.1 | ADK94697.1 | *Plutella xylostella* | Arthropoda | Insecta | Yes | DEGSLFEVKSTAGD | |
| 4 | | HM370510.1 | ADK94698.1 | *Plutella xylostella* | Arthropoda | Insecta | Yes | DEGSLFEVKSTAGD | |
| 4 | | HM370511.1 | ADK94699.1 | *Plutella xylostella* | Arthropoda | Insecta | Yes | DEGSLFEVKSTAGD | |
| 4 | | HM593518.1 | ADP37711.1 | *Helicoverpa armigera* | Arthropoda | Insecta | Yes | DEGSLFEVRSTAGD | |
| 4 | | HM769899.1 | ADL27420.1 | *Chironomus riparius* | Arthropoda | Insecta | Yes | DEGSLFEVRSTAGD | |
| 4 | | HQ012004.2 | ADV03160.1 | *Spodoptera litura* | Arthropoda | Insecta | Yes | DEGSLFEVRATAGD | |
| 4 | | HQ107971.1 | ADV58254.1 | *Plutella xylostella* | Arthropoda | Insecta | Yes | DEGSLFEVKSTAGD | |
| 4 | | HQ107972.1 | ADV58255.1 | *Plutella xylostella* | Arthropoda | Insecta | Yes | DEGSLFEVKSTAGD | |
| 4 | | JF421286.1 | AEB52075.1 | *Microdera punctipennis* | Arthropoda | Insecta | Yes | DEGSLFEVRATAGD | |
| 4 | | JF836795.1 | AEI58997.1 | *Bombyx mori* | Arthropoda | Insecta | Yes | DEGSLFEVKATAGD | |
| 4 | | JN393894.1 | AEX63624.1 | *Aedes aegypti* | Arthropoda | Insecta | - | – | Partial sequence without the Ser-containing region; Not used included in |
| 4 | | JN863694.2 | AFK93489.2 | *Cydia pomonella* | Arthropoda | Insecta | Yes | DEGSLFEVKATAGD | |
| 4 | | JN863695.1 | AFK93490.1 | *Cydia pomonella* | Arthropoda | Insecta | Yes | DEGSLFEVKATAGD | |
| 4 | | JN971028.1 | AFC76151.1 | *Quadrastichus erythrinae* | Arthropoda | Insecta | Yes | DEGSLFEVKATAGD | |
| 4 | | JQ219849.1 | AFE88580.1 | *Tenebrio molitor* | Arthropoda | Insecta | Yes | DEGSLFEVRATAGD | |
| 4 | | JQ316541.2 | AFO70209.1 | *Hypena tristalis* | Arthropoda | Insecta | Yes | DEGSLFEVRSTAGD | |
| 4 | | JQ655741.1 | AFX84617.1 | *Frankliniella occidentalis* | Arthropoda | Insecta | Yes | DEGSLFEVKSTAGD | |
| 4 | | JQ839279.1 | AFM45297.1 | *Habrobracon hebetor* | Arthropoda | Insecta | Yes | DEGSLFEVKSTAGD | |
| 4 | | JQ859844.2 | AFN08643.1 | *Oxya chinensis* | Arthropoda | Insecta | Yes | SEGSLFEVKATAGD | |
| 4 | | JX088377.1 | AGF34717.1 | *Cotesia vestalis* | Arthropoda | Insecta | Yes | DEGSLFEVKSTAGD | |
| 4 | | JX627810.1 | AFX84560.1 | *Lygus hesperus* | Arthropoda | Insecta | Yes | DEGSLFEVRATAGD | |
| 4 | | JX678981.1 | AFU06382.1 | *Grapholita molesta* | Arthropoda | Insecta | Yes | DEGSLFEVKATAGD | |
| 4 | | JX680669.1 | AGR84216.1 | *Melitaea cinxia* | Arthropoda | Insecta | Yes | DEGSLFEVKATAGD | |
| 4 | | JX680670.1 | AGR84217.1 | *Melitaea cinxia* | Arthropoda | Insecta | Yes | DEGSLFEVKATAGD | |
| 4 | | JX680671.1 | AGR84218.1 | *Melitaea cinxia* | Arthropoda | Insecta | Yes | DEGSLFEVKATAGD | |
| 4 | | JX680672.1 | AGR84219.1 | *Melitaea cinxia* | Arthropoda | Insecta | Yes | DEGSLFEVKATAGD | |
| 4 | | JX680675.1 | AGR84222.1 | *Melitaea cinxia* | Arthropoda | Insecta | Yes | DEGSLFEVRSTAGD | |
| 4 | | JX680676.1 | AGR84223.1 | *Melitaea cinxia* | Arthropoda | Insecta | Yes | DEGSLFEVKATAGD | |
| 4 | | JX680677.1 | AGR84224.1 | *Melitaea cinxia* | Arthropoda | Insecta | Yes | DEGSLFEVKATAGD | |

| | | | | | | | |
|---|---|---|---|---|---|---|---|
| 4 | JX680680.1 | AGR84227.1 | *Melitaea cinxia* | Arthropoda | Insecta | Yes | SEGSLFEVRSTAGD |
| 4 | JX961639.1 | AGN32394.1 | *Bactrocera correcta* | Arthropoda | Insecta | Yes | DEGSLFEVRATAGD |
| 4 | KA646595.1 | AFP61224.1 | *Musca domestica* | Arthropoda | Insecta | Yes | DEGSLFEVRSTAGD |
| 4 | KC161300.1 | AGE92595.1 | *Ericerus pela* | Arthropoda | Insecta | Yes | SEGSLFEVLSTAGD |
| 4 | KC544270.1 | AHA36970.1 | *Leptinotarsa decemlineata* | Arthropoda | Insecta | Yes | DEGSLFEVKSTAGD |
| 4 | KC620440.1 | AHE77387.1 | *Lissorhoptrus oryzophilus* | Arthropoda | Insecta | Yes | DEGSLFEVRSTAGD |
| 4 | KC860254.1 | AGQ45967.1 | *Diamesa cinerella* | Arthropoda | Insecta | Yes | DQGSLFEVKSTAGD |
| 4 | KF018929.1 | AGT99186.1 | *Corythucha ciliata* | Arthropoda | Insecta | Yes | DEGSLFEVKSTAGD |
| 4 | KF214746.1 | AHF52926.1 | *Colaphellus bowringi* | Arthropoda | Insecta | Yes | DEGSLFEVKSTAGD |
| 4 | KF730249.1 | AIM18802.1 | *Empoasca onukii* | Arthropoda | Insecta | Yes | SEGSLFEVKSTAGD |
| 4 | KF731995.1 | AHF81959.1 | *Leguminivora glycinivorella* | Arthropoda | Insecta | No | -EDGIFEVKSTAGD |
| 4 | KF771049.1 | AHG94986.1 | *Aphis glycines* | Arthropoda | Insecta | Yes | DEGSIFEVKSTAGD |
| 4 | KF792067.1 | AHH25012.1 | *Agasicles hygrophila* | Arthropoda | Insecta | Yes | DEGSLFEVKATAGD |
| 4 | KJ541738.1 | AIA62361.1 | *Bactrocera minax* | Arthropoda | Insecta | Yes | DEGSLFEVRATAGD |
| 4 | KJ701421.1 | AID61521.1 | *Pieris rapae rapae* | Arthropoda | Insecta | Yes | DEGSLFEVKATAGD |
| 4 | KJ813013.1 | AIS72815.1 | *Sitodiplosis mosellana* | Arthropoda | Insecta | Yes | DDGSLFEVKSTAGD |
| 4 | KJ867518.1 | AIK01869.1 | *Orius sauteri* | Arthropoda | Insecta | Yes | DEGSLFEVRATAGD |
| 4 | KJ909505.1 | AIL52739.1 | *Phenacoccus solenopsis* | Arthropoda | Insecta | Yes | DEGSLFEVKSTAGD |
| 4 | KM103515.1 | AJZ68816.1 | *Cotesia chilonis* | Arthropoda | Insecta | Yes | DEGSLFEVKSTAGD |
| 4 | KM112021.1 | AJZ68832.1 | *Zeugodacus cucurbitae* | Arthropoda | Insecta | Yes | DEGSLFEVRATAGD |
| 4 | KT003964.1 | ANJ86404.1 | *Grapholita molesta* | Arthropoda | Insecta | Yes | DEGSLFEVKATAGD |
| 4 | KT225460.1 | ALL42052.1 | *Antheraea pernyi* | Arthropoda | Insecta | Yes | DEGSLFEVRATAGD |
| 4 | KU050681.1 | ANA11230.1 | *Dastarcus helophoroides* | Arthropoda | Insecta | Yes | DEGSLFEVKSTAGD |
| 4 | KU050682.1 | ANA11231.1 | *Dastarcus helophoroides* | Arthropoda | Insecta | Yes | DEGSLFEVKATAGD |
| 4 | KU159184.1 | AMX23517.1 | *Monochamus alternatus* | Arthropoda | Insecta | Yes | DEGSLFEVKATAGD |
| 4 | KU516012.1 | AMK38874.1 | *Colaphellus bowringi* | Arthropoda | Insecta | Yes | DEGSIFEVKATAGD |
| 4 | KX231836.1 | ANM71228.1 | *Sitophilus oryzae* | Arthropoda | Insecta | Yes | DEGSLFEVKATAGD |
| 4 | KY231148.1 | ARQ84028.1 | *Liriomyza trifolii* | Arthropoda | Insecta | Yes | DEGSIFEVKATAGD |
| 4 | KY457331.1 | ASA47799.1 | *Hermetia illucens* | Arthropoda | Insecta | Yes | DEGSLFEVKSTAGD |
| 4 | KY695168.1 | AVR54974.1 | *Sitophilus oryzae* | Arthropoda | Insecta | Yes | DEGSLFEVKSTAGD |
| 4 | KY705383.1 | ARW29609.1 | *Corythucha ciliata* | Arthropoda | Insecta | Yes | DEGSLFEVKSTAGD |
| 4 | KY705384.1 | ARW29610.1 | *Corythucha ciliata* | Arthropoda | Insecta | Yes | DEGSIFEVKATAGD |
| 4 | KY705385.1 | ARW29611.1 | *Corythucha ciliata* | Arthropoda | Insecta | Yes | DEGSIFEVKATAGD |
| 4 | KY930331.1 | ATL76715.1 | *Eogystia hippophaecolus* | Arthropoda | Insecta | Yes | DEGSLFEVKSTAGD |
| 4 | KY933451.1 | ARQ84041.1 | *Liriomyza trifolii* | Arthropoda | Insecta | Yes | DEGSLFEVRSTAGD |
| 4 | KY933452.1 | ARQ84042.1 | *Liriomyza trifolii* | Arthropoda | insecta | Yes | DEGSLFEVKSTAGD |
| 4 | MF773976.1 | AXB26576.1 | *Anaphothrips obscurus* | Arthropoda | Insecta | Yes | DEGSLFEVRSTAGD |
| 4 | MG387117.1 | AWS20697.1 | *Aphidius gifuensis* | Arthropoda | Insecta | Yes | DEGSLFEVRSTAGD |
| 4 | MH042023.1 | AXJ21598.1 | *Mythimna separata* | Arthropoda | Insecta | Yes | DEGSIFEVKATAGD |
| 4 | MH366003.1 | QBB01812.1 | *Cotesia chilonis* | Arthropoda | Insecta | Yes | DEGSLFEVKSTAGD |
| 4 | MK000439.1 | QCI56580.1 | *Tribolium castaneum* | Arthropoda | Insecta | Yes | DEGSIFEVKATAGD |
| 4 | MK000440.1 | QCI56581.1 | *Tribolium castaneum* | Arthropoda | Insecta | Yes | DEGSIFEVKATAGD |
| 4 | MK168341.1 | QAV55738.1 | *Sogatella furcifera* | Arthropoda | Insecta | Yes | DEGSLFEVRSTAGD |
| 4 | MN138034.1 | QEL52753.1 | *Agasicles hygrophila* | Arthropoda | Insecta | Yes | DEGSLFEVKATAGD |
| 4 | MN190009.1 | QMU24016.1 | *Bemisia tabaci* | Arthropoda | Insecta | Yes | DEGSLFEVRATAGD |
| 4 | MN190010.1 | QMU24017.1 | *Bemisia tabaci* | Arthropoda | Insecta | Yes | DEGSLFEVRATAGD |
| 4 | MN190012.1 | QMU24019.1 | *Bemisia tabaci* | Arthropoda | Insecta | Yes | DEGSLFEVRATAGD |
| 4 | MN190014.1 | QMU24021.1 | *Bemisia tabaci* | Arthropoda | Insecta | Yes | DEGSLFEVRATAGD |
| 4 | MN190015.1 | QMU24022.1 | *Bemisia tabaci* | Arthropoda | Insecta | Yes | DEGSLFEVRATAGD |
| 4 | MN190016.1 | QMU24023.1 | *Bemisia tabaci* | Arthropoda | Insecta | Yes | DEGSLFEVRATAGD |
| 4 | MN190017.1 | QMU24024.1 | *Bemisia tabaci* | Arthropoda | Insecta | Yes | DEGSLFEVRATAGD |
| 4 | MN190018.1 | QMU24025.1 | *Bemisia tabaci* | Arthropoda | Insecta | Yes | DEGSLFEVKSTAGD |
| 4 | MN241441.1 | QOY58048.1 | *Harmonia axyridis* | Arthropoda | Insecta | Yes | DEGSLFEVKSTAGD |
| 4 | MN480721.1 | QGA73357.1 | *Spodoptera frugiperda* | Arthropoda | Insecta | Yes | DEGSLFEVRATAGD |
| 4 | MN895063.1 | QTA73201.1 | *Monochamus alternatus* | Arthropoda | Insecta | Yes | DEGSLFEVRATAGD |
| 4 | MN895064.1 | QTA73202.1 | *Monochamus alternatus* | Arthropoda | Insecta | Yes | DEGSLFEVRSTAGD |
| 4 | MT157251.1 | QUS47835.1 | *Macrosiphum euphorbiae* | Arthropoda | Insecta | Yes | DEGSIFEVKSTAGD |
| 4 | MT261583.1 | QSL97649.1 | *Hyphantria cunea* | Arthropoda | Insecta | Yes | AEGSLFEVKATSGD |
| 4 | MW321609.1 | QWV59537.1 | *Lasioderma serricorne* | Arthropoda | Insecta | Yes | SEGSLFEVRATAGD |
| 4 | MW321612.1 | QWV59540.1 | *Lasioderma serricorne* | Arthropoda | Insecta | Yes | NEGSLFEVRSTAGD |
| 4 | MW691114.1 | QWV59541.1 | *Pieris rapae* | Arthropoda | Insecta | Yes | DEGSLFEVKATAGD |
| 4 | MW691115.1 | QWV59542.1 | *Pieris rapae* | Arthropoda | Insecta | Yes | DEGSLFEVKATAGD |
| 4 | MW691117.1 | QWV59544.1 | *Pieris rapae* | Arthropoda | Insecta | Yes | DEGSLFEVKATAGD |
| 4 | NM_001043931.1 | NP_001037396.1 | *Bombyx mori* | Arthropoda | Insecta | Yes | DEGSLFEVKSTAGD |
| 4 | NM_001309597.1 | NP_001296526.1 | *Bombyx mori* | Arthropoda | Insecta | Yes | DEGSLFEVKSTAGD |
| 4 | NM_079615.4 | NP_524339.1 | *Drosophila melanogaster* | Arthropoda | Insecta | Yes | DEGSLFEVKATAGD |
| 4 | NM_080188.3 | NP_524927.2 | *Drosophila melanogaster* | Arthropoda | Insecta | Yes | DEGSLFEVRSTAGD |
| 4 | NM_169441.2 | NP_731651.1 | *Drosophila melanogaster* | Arthropoda | Insecta | Yes | DEGSLFEVRSTAGD |
| 4 | NM_169469.2 | NP_731716.1 | *Drosophila melanogaster* | Arthropoda | Insecta | Yes | DEGSLFEVRSTAGD |
| 4 | NM_176486.2 | NP_788663.1 | *Drosophila melanogaster* | Arthropoda | Insecta | Yes | DEGSLFEVRSTAGD |
| 4 | X78403.1 | CAA55168.1 | *Drosophila auraria* | Arthropoda | Insecta | Yes | DEGSLFEVRATAGD |
| 4 | XM_001864875.2 | XP_001864910.2 | *Culex quinquefasciatus* | Arthropoda | Insecta | Yes | DEGSLFEVKATAGD |
| 4 | XM_001945751.5 | XP_001945786.2 | *Acyrthosiphon pisum* | Arthropoda | Insecta | Yes | DEGSIFEVKSTAGD |
| 4 | XM_001949802.5 | XP_001949837.1 | *Acyrthosiphon pisum* | Arthropoda | Insecta | Yes | DEGSLFEVKSTAGD |
| 4 | XM_002424282.1 | XP_002424327.1 | *Pediculus humanus corporis* | Arthropoda | Insecta | Yes | DEGSLFEVRATAGD |
| 4 | XM_004536177.4 | XP_004536234.1 | *Ceratitis capitata* | Arthropoda | Insecta | Yes | DEGSLFEVRATAGD |
| 4 | XM_005190957.3 | XP_005191014.3 | *Musca domestica* | Arthropoda | Insecta | Yes | DEGSLFEVRSTAGD |
| 4 | XM_008478797.3 | XP_008477019.1 | *Diaphorina citri* | Arthropoda | Insecta | No | DEGALFEVRSTAGD |
| 4 | XM_008478798.3 | XP_008477020.1 | *Diaphorina citri* | Arthropoda | Insecta | No | DEGALFEVRSTAGD |
| 4 | XM_008478800.2 | XP_008477022.1 | *Diaphorina citri* | Arthropoda | Insecta | No | DEGALFEVRSTAGD |
| 4 | XM_011316813.1 | XP_011315115.1 | *Fopius arisanus* | Arthropoda | Insecta | Yes | DEGSLFEVRSTAGD |
| 4 | XM_011550936.2 | XP_011549238.2 | *Plutella xylostella* | Arthropoda | Insecta | Yes | DEGSLFEVKSTAGD |
| 4 | XM_012397804.2 | XP_012253227.1 | *Athalia rosae* | Arthropoda | Insecta | Yes | DEGSLFEVRATAGD |
| 4 | XM_013255123.1 | XP_013110577.1 | *Stomoxys calcitrans* | Arthropoda | Insecta | Yes | DEGSLFEVRATAGD |
| 4 | XM_014754254.1 | XP_014609740.1 | *Polistes canadensis* | Arthropoda | Insecta | Yes | DEGSLFEVKATAGD |
| 4 | XM_015271680.1 | XP_015127166.1 | *Diachasma alloeum* | Arthropoda | Insecta | Yes | DEGSLFEVKSTAGD |
| 4 | XM_015511172.1 | XP_015366658.1 | *Diuraphis noxia* | Arthropoda | Insecta | Yes | DEGSIFEVKSTAGD |
| 4 | XM_015516481.1 | XP_015371967.1 | *Diuraphis noxia* | Arthropoda | Insecta | Yes | DEGSLFEVKSTAGD |
| 4 | XM_015516573.1 | XP_015372059.1 | *Diuraphis noxia* | Arthropoda | Insecta | Yes | DEGSIFEVKSTAGD |
| 4 | XM_015522949.1 | XP_015378435.1 | *Diuraphis noxia* | Arthropoda | Insecta | Yes | DEGSIFEVKSTAGD |
| 4 | XM_015659127.1 | XP_015514613.1 | *Neodiprion lecontei* | Arthropoda | Insecta | Yes | DEGSLFEVKATAGD |
| 4 | XM_015977915.1 | XP_015833401.1 | *Tribolium castaneum* | Arthropoda | Insecta | No | -ADGIFEVKSTAGD |
| 4 | XM_015977916.1 | XP_015833402.1 | *Tribolium castaneum* | Arthropoda | Insecta | No | AYGLDKKVKSTAGD |
| 4 | XM_017637032.1 | XP_017492521.1 | *Rhagoletis zephyria* | Arthropoda | Insecta | Yes | DEGSLFEVRATAGD |
| 4 | XM_017931378.1 | XP_017786867.1 | *Nicrophorus vespilloides* | Arthropoda | Insecta | Yes | DEGSLFEVRSTAGD |
| 4 | XM_018481181.2 | XP_018336683.1 | *Agrilus planipennis* | Arthropoda | Insecta | Yes | DEGSLFEVKATAGD |
| 4 | XM_018705664.1 | XP_018561180.1 | *Anoplophora glabripennis* | Arthropoda | Insecta | Yes | DEGSLFEVRSTAGD |
| 4 | XM_019672735.2 | XP_019528280.2 | *Aedes albopictus* | Arthropoda | Insecta | Yes | DEGSIFEVKATAGD |
| 4 | XM_019672747.2 | XP_019528292.2 | *Aedes albopictus* | Arthropoda | Insecta | Yes | DEGSIFEVKATAGD |
| 4 | XM_019698586.2 | XP_019554131.2 | *Aedes albopictus* | Arthropoda | Insecta | Yes | DEGSIFEVKATAGD |
| 4 | XM_019698588.2 | XP_019554133.2 | *Aedes albopictus* | Arthropoda | Insecta | Yes | DEGSLFEVKATAGD |
| 4 | XM_019698591.2 | XP_019554136.2 | *Aedes albopictus* | Arthropoda | Insecta | Yes | DEGSLFEVRATAGD |
| 4 | XM_019698592.2 | XP_019554137.2 | *Aedes albopictus* | Arthropoda | Insecta | Yes | DEGSLFEVRATAGD |
| 4 | XM_019905100.1 | XP_019760659.1 | *Dendroctonus Ponderosae* | Arthropoda | Insecta | Yes | DEGSLFEVRSTAGD |
| 4 | XM_019905106.1 | XP_019760665.1 | *Dendroctonus Ponderosae* | Arthropoda | Insecta | Yes | DEGSLFEVRATAGD |
| 4 | XM_020039259.1 | XP_019894818.1 | *Musca Domestica* | Arthropoda | Insecta | Yes | DEGSLFEVRSTAGD |
| 4 | XM_021333392.1 | XP_021189067.1 | *Helicoverpa armigera* | Arthropoda | Insecta | Yes | AEGSLFEVKSTAGD |
| 4 | XM_021837957.1 | XP_021693649.1 | *Aedes aegypti* | Arthropoda | Insecta | Yes | DEGSLFEVRATAGD |
| 4 | XM_021837961.1 | XP_021693653.1 | *Aedes aegypti* | Arthropoda | Insecta | Yes | DQGSLFEVRATAGD |
| 4 | XM_021837962.1 | XP_021693654.1 | *Aedes Aegypti* | Arthropoda | Insecta | Yes | DQGSLFEVRATAGD |
| 4 | XM_021837963.1 | XP_021693655.1 | *Aedes Aegypti* | Arthropoda | Insecta | Yes | DEGSLFEVKSTAGD |
| 4 | XM_021857641.1 | XP_021713333.1 | *Aedes Aegypti* | Arthropoda | Insecta | Yes | DEGSLFEVRATAGD |
| 4 | XM_022305334.1 | XP_022161026.1 | *Myzus persicae* | Arthropoda | Insecta | Yes | DEGSLFEVKSTAGD |
| 4 | XM_022312213.1 | XP_022167905.1 | *Myzus persicae* | Arthropoda | Insecta | Yes | DEGSIFEVKSTAGD |
| 4 | XM_022327049.1 | XP_022182741.1 | *Myzus persicae* | Arthropoda | Insecta | Yes | DEGSIFEVKSTAGD |
| 4 | XM_023054585.1 | XP_022910353.1 | *Onthophagus taurus* | Arthropoda | Insecta | Yes | DEGSIFEVKATAGD |
| 4 | XM_023443509.1 | XP_023299277.1 | *Lucilia cuprina* | Arthropoda | Insecta | Yes | DEGSIFEVKATAGD |
| 4 | XM_023869487.2 | XP_023725255.1 | *Cryptotermes secundus* | Arthropoda | Insecta | Yes | DEGSIFEVKATAGD |
| 4 | XM_025334484.1 | XP_025190269.1 | *Melanaphis sacchari* | Arthropoda | Insecta | Yes | DEGSLFEVRSTAGD |

| | | | | | | | | | |
|---|---|---|---|---|---|---|---|---|---|
| 4 | | XM_025342129.1 | XP_025197914.1 | *Melanaphis sacchari* | Arthropoda | Insecta | Yes | DEGSIFEVKATAGD | |
| 4 | | XM_025343133.1 | XP_025198918.1 | *Melanaphis sacchari* | Arthropoda | Insecta | Yes | DEGSIFEVKATAGD | |
| 4 | | XM_025344042.1 | XP_025199827.1 | *Melanaphis sacchari* | Arthropoda | Insecta | Yes | DEGSIFEVKATAGD | |
| 4 | | XM_025345168.1 | XP_025200953.1 | *Melanaphis sacchari* | Arthropoda | Insecta | Yes | DEGSLFEVRSTAGD | |
| 4 | | XM_025346084.1 | XP_025201869.1 | *Melanaphis sacchari* | Arthropoda | Insecta | Yes | DEGSLFEVRSTAGD | |
| 4 | | XM_025346922.1 | XP_025202707.1 | *Melanaphis sacchari* | Arthropoda | Insecta | Yes | DEGSLFEVRSTAGD | |
| 4 | | XM_025350710.1 | XP_025206495.1 | *Melanaphis sacchari* | Arthropoda | Insecta | Yes | DEGSLFEVRSTAGD | |
| 4 | | XM_025350978.1 | XP_025206763.1 | *Melanaphis sacchari* | Arthropoda | Insecta | Yes | DEGSLFEVRSTAGD | |
| 4 | | XM_025549051.1 | XP_025404836.1 | *Sipha flava* | Arthropoda | Insecta | Yes | DEGSLFEVRSTAGD | |
| 4 | | XM_025549993.1 | XP_025405778.1 | *Sipha flava* | Arthropoda | Insecta | Yes | DEGSLFEVRSTAGD | |
| 4 | | XM_025558496.1 | XP_025414281.1 | *Sipha flava* | Arthropoda | Insecta | Yes | DEGSLFEVRSTAGD | |
| 4 | | XM_026624612.1 | XP_026480397.1 | *Ctenocephalides felis* | Arthropoda | Insecta | Yes | DEGSIFEVKATAGD | |
| 4 | | XM_026624613.1 | XP_026480398.1 | *Ctenocephalides felis* | Arthropoda | Insecta | Yes | DEGSIFEVKATAGD | |
| 4 | | XM_026883810.1 | XP_026739611.1 | *Trichoplusia ni* | Arthropoda | Insecta | Yes | DEGSIFEVKATAGD | |
| 4 | | XM_026883995.1 | XP_026739796.1 | *Trichoplusia ni* | Arthropoda | Insecta | Yes | DEGSIFEVKATAGD | |
| 4 | | XM_026958889.1 | XP_026814690.1 | *Rhopalosiphum maidis* | Arthropoda | Insecta | Yes | DEGSLFEVRSTAGD | |
| 4 | | XM_026959488.1 | XP_026815289.1 | *Rhopalosiphum maidis* | Arthropoda | Insecta | Yes | DEGSLFEVRSTAGD | |
| 4 | | XM_026964118.2 | XP_026819919.1 | *Rhopalosiphum maidis* | Arthropoda | Insecta | Yes | DEGSLFEVKSTAGD | |
| 4 | | XM_026964932.1 | XP_026820733.1 | *Rhopalosiphum maidis* | Arthropoda | Insecta | Yes | DEGSIFEVKSTAGD | |
| 4 | | XM_027981156.1 | XP_027836957.1 | *Aphis gossypii* | Arthropoda | Insecta | Yes | DEGSIFEVKSTAGD | |
| 4 | | XM_027984663.1 | XP_027840464.1 | *Aphis gossypii* | Arthropoda | Insecta | No | DEGLIFEVKSTAGD | |
| 4 | | XM_027987439.1 | XP_027843240.1 | *Aphis gossypii* | Arthropoda | Insecta | Yes | DEGSIFEVKSTAGD | |
| 4 | | XM_027997432.1 | XP_027853233.1 | *Aphis gossypii* | Arthropoda | Insecta | Yes | DEGSIFEVKSTAGD | |
| 4 | | XM_027998713.1 | XP_027854514.1 | *Aphis gossypii* | Arthropoda | Insecta | Yes | DEGSIFEVKSTAGD | |
| 4 | | XM_028273705.1 | XP_028129506.1 | *Diabrotica virgifera virgifera* | Arthropoda | Insecta | Yes | DEGSLFEVRSTAGD | |
| 4 | | XM_028273710.1 | XP_028129511.1 | *Diabrotica virgifera virgifera* | Arthropoda | Insecta | Yes | DEGSLFEVRSTAGD | |
| 4 | | XM_028278425.1 | XP_028134226.1 | *Diabrotica virgifera virgifera* | Arthropoda | Insecta | Yes | DEGSLFEVRSTAGD | |
| 4 | | XM_028296548.1 | XP_028152349.1 | *Diabrotica virgifera virgifera* | Arthropoda | Insecta | - | – | Partial sequence without the Ser-containing region; Not used included in [illegible] |
| 4 | | XM_029867428.1 | XP_029723288.1 | *Aedes albopictus* | Arthropoda | Insecta | Yes | DEGSLFEVRATAGD | |
| 4 | | XM_029867429.1 | XP_029723289.1 | *Aedes albopictus* | Arthropoda | Insecta | Yes | DEGSLFEVRATAGD | |
| 4 | | XM_029867430.1 | XP_029723290.1 | *Aedes albopictus* | Arthropoda | Insecta | Yes | DEGSLFEVRATAGD | |
| 4 | | XM_029867446.1 | XP_029723306.1 | *Aedes albopictus* | Arthropoda | Insecta | Yes | DEGSLFEVRATAGD | |
| 4 | | XM_029867448.1 | XP_029723308.1 | *Aedes albopictus* | Arthropoda | Insecta | Yes | DQGSLFEVKATAGD | |
| 4 | | XM_029869301.1 | XP_029725161.1 | *Aedes albopictus* | Arthropoda | Insecta | Yes | DEGSLFEVKATAGD | |
| 4 | | XM_029869302.1 | XP_029725162.1 | *Aedes albopictus* | Arthropoda | Insecta | Yes | DEGSLFEVRATAGD | |
| 4 | | XM_029872754.1 | XP_029728614.1 | *Aedes albopictus* | Arthropoda | Insecta | Yes | DEGSLFEVRATAGD | |
| 4 | | XM_029872755.1 | XP_029728615.1 | *Aedes albopictus* | Arthropoda | Insecta | Yes | DEGSLFEVRATAGD | |
| 4 | | XM_029872756.1 | XP_029728616.1 | *Aedes albopictus* | Arthropoda | Insecta | Yes | DEGSLFEVKATAGD | |
| 4 | | XM_029872757.1 | XP_029728617.1 | *Aedes albopictus* | Arthropoda | Insecta | Yes | DQGSLFEVKATAGD | |
| 4 | | XM_029876940.1 | XP_029732800.1 | *Aedes albopictus* | Arthropoda | Insecta | Yes | DEGSLFEVKATAGD | |
| 4 | | XM_031483209.1 | XP_031339069.1 | *Aedes albopictus* | Arthropoda | Insecta | Yes | DEGSLFEVRSTAGD | |
| 4 | | XM_031483210.1 | XP_031339070.1 | *Photinus pyralis* | Arthropoda | Insecta | Yes | DEGSLFEVKSTAGD | |
| 4 | | XM_031483367.1 | XP_031339227.1 | *Photinus pyralis* | Arthropoda | Insecta | Yes | DEGSLFEVRSTAGD | |
| 4 | | XM_031483368.1 | XP_031339228.1 | *Photinus pyralis* | Arthropoda | Insecta | Yes | DEGSLFEVRSTAGD | |
| 4 | | XM_031502141.1 | XP_031358001.1 | *Photinus pyralis* | Arthropoda | Insecta | Yes | DEGSLFEVKSTAGD | |
| 4 | | XM_031782483.1 | XP_031638343.1 | *Photinus pyralis* | Arthropoda | Insecta | Yes | DEGSLFEVRSTAGD | |
| 4 | | XM_031782484.1 | XP_031638344.1 | *Contarinia nasturtii* | Arthropoda | Insecta | Yes | DEGSLFEVRSTAGD | |
| 4 | | XM_033372596.1 | XP_033228487.1 | *Amblyraja radiata* | Arthropoda | Insecta | Yes | DEGSLFEVKSTAGD | |
| 4 | | XM_034401249.1 | XP_034257140.1 | *Thrips palmi* | Arthropoda | Insecta | Yes | DEGSLFEVRSTAGD | |
| 4 | | XM_035083965.1 | XP_034939856.1 | *Chelonus insularis* | Arthropoda | Insecta | Yes | DEGSLFEVKATAGD | |
| 4 | | XM_035916939.1 | XP_035772832.1 | *Anopheles albimanus* | Arthropoda | Insecta | Yes | DEGSLFEVRSTAGD | |
| 4 | | XM_035940236.1 | XP_035796129.1 | *Anopheles albimanus* | Arthropoda | Insecta | Yes | DEGSLFEVRATAGD | |
| 4 | | XM_036046027.1 | XP_035901920.1 | *Anopheles stephensi* | Arthropoda | Insecta | Yes | DEGSLFEVRATAGD | |
| 4 | | XM_036046029.1 | XP_035901922.1 | *Anopheles stephensi* | Arthropoda | Insecta | Yes | DEGSLFEVRATAGD | |
| 4 | | XM_036046030.1 | XP_035901923.1 | *Anopheles stephensi* | Arthropoda | Insecta | Yes | DEGSLFEVRATAGD | |
| 4 | | XM_036472402.1 | XP_036328295.1 | *Rhagoletid pomonella* | Arthropoda | Insecta | Yes | DEGSLFEVRATAGD | |
| 4 | | XM_037175227.1 | XP_037031122.1 | *Bradysia coprophila* | Arthropoda | Insecta | Yes | DEGSLFEVKSTAGD | |
| 4 | | XM_037175228.1 | XP_037031123.1 | *Bradysia coprophila* | Arthropoda | Insecta | Yes | DEGSIFEVKATAGD | |
| 4 | | XM_037177791.1 | XP_037033686.1 | *Bradysia coprophila* | Arthropoda | Insecta | Yes | DEGSIFEVKATAGD | |
| 4 | | XM_037960142.1 | XP_037816070.1 | *Lucilia sericata* | Arthropoda | Insecta | Yes | DEGSIFEVKATAGD | |
| 4 | | XM_038020963.1 | XP_037876891.1 | *Bombyx mori* | Arthropoda | Insecta | Yes | DEGSLFEVKSTAGD | |
| 4 | | XM_038110358.1 | XP_037966286.1 | *Plutella xylostella* | Arthropoda | Insecta | Yes | DEGSIFEVKATAGD | |
| 4 | | XM_039585513.1 | XP_039441447.1 | *Culex pipiens pallens* | Arthropoda | Insecta | Yes | DEGSLFEVRSTAGD | |
| 4 | | XM_039597734.1 | XP_039453668.1 | *Culex pipiens pallens* | Arthropoda | Insecta | Yes | DEGSLFEVRSTAGD | |
| 4 | | XM_040093844.1 | XP_039949778.1 | *Bactrocera tryoni* | Arthropoda | Insecta | Yes | DEGSIFEVKATAGD | |
| 4 | | XM_040298425.1 | XP_040154359.1 | *Anopheles arabiensis* | Arthropoda | Insecta | Yes | DEGSIFEVKATAGD | |
| 4 | | XM_040298618.1 | XP_040154552.1 | *Anopheles arabiensis* | Arthropoda | Insecta | Yes | DEGSIFEVKATAGD | |
| 4 | | XM_040298706.1 | XP_040154640.1 | *Anopheles arabiensis* | Arthropoda | Insecta | Yes | DEGSLFEVRATAGD | |
| 4 | | XM_040369812.1 | XP_040225746.1 | *Anopheles coluzzii* | Arthropoda | Insecta | Yes | DEGSLFEVRATAGD | |
| 4 | | XM_040370239.1 | XP_040226173.1 | *Anopheles coluzzii* | Arthropoda | Insecta | Yes | DEGSLFEVRATAGD | |
| 4 | | XM_041907784.1 | XP_041763718.1 | *Anopheles merus* | Arthropoda | Insecta | Yes | DEGSLFEVRATAGD | |
| 4 | | XM_041907928.1 | XP_041763862.1 | *Anopheles merus* | Arthropoda | Insecta | Yes | DEGSLFEVRATAGD | |
| 4 | | XM_041908012.1 | XP_041763946.1 | *Anopheles merus* | Arthropoda | Insecta | Yes | DEGSLFEVRATAGD | |
| 4 | | XM_044408958.1 | XP_044264893.1 | *Tribolium madens* | Arthropoda | Insecta | Yes | DEGSLFEVRATAGD | |
| 4 | | XM_044409295.1 | XP_044265230.1 | *Tribolium madens* | Arthropoda | Insecta | Yes | DEGSLFEVRATAGD | |
| 4 | | XM_044409722.1 | XP_044265657.1 | *Tribolium madens* | Arthropoda | Insecta | Yes | DEGSLFEVKATAGD | |
| 4 | | XM_969349.3 | XP_974442.1 | *Tribolium castaneum* | Arthropoda | Insecta | Yes | DEGSLFEVKATAGD | |
| 4 | HSP70cA1 | XM_013527000.2 | XP_013382454.1 | *Lingula anatina* | Brachiopoda | Lingulata | Yes | ADGSLFEVRSTAGD | |
| 4 | HSP70cA1 | GSADVT00001811001_v2.0 | - | *Adineta vaga* | Rotifera | Eurotatoria | Yes | AGGSVFEVRSTAGD | |
| 4 | HSP70cA2 | GSADVT00016830001_v2.0 | - | *Adineta vaga* | Rotifera | Eurotatoria | Yes | VSGSVFEVKATAGD | |
| 4 | | HQ693698.2 | ADV57677.1 | *Eisenia fetida* | Annelida | Clitellata | No | -EDGIFEVKSTSGD | |
| 4 | HSP70cA1i | EF580992.1 | ABU63808.1 | *Paralvinella grasslei* | Annelida | Polychaeta | Yes | ADGSLFEVKSTAGD | |
| 4 | | JX560964.1 | AGH18393.1 | *Alvinella pompejana* | Annelida | Polychaeta | Yes | GDGSLFEVKSTAGD | |
| 4 | | AB122063.1 | BAD15286.1 | *Crassostrea gigas* | Mollusca | Bivalvia | Yes | DEGSIFEVRSTAGD | |
| 4 | | AB180908.1 | BAD99026.1 | *Mytilus galloprovincialis* | Mollusca | Bivalvia | Yes | DEGSLFEVRSTAGD | |
| 4 | | AB180909.1 | BAD99027.1 | *Mytilus galloprovincialis* | Mollusca | Bivalvia | Yes | DEGSLFEVRSTAGD | |
| 4 | | AB549340.1 | BAJ83619.1 | *Crassostrea gigas* | Mollusca | Bivalvia | Yes | DEGSIFEVRSTAGD | |
| 4 | | AF416608.1 | AAM46634.1 | *Ostrea edulis* | Mollusca | Bivalvia | Yes | DEGSIFEVLSTAGD | |
| 4 | | AF416609.1 | AAM46635.1 | *Ostrea edulis* | Mollusca | Bivalvia | Yes | DEGSIFEVRSTAGD | |
| 4 | | AJ271444.1 | CAB89802.1 | *Crassostrea virginica* | Mollusca | Bivalvia | Yes | DEGSIFEVRSTAGD | |
| 4 | | FJ157365.1 | ACH95805.1 | *Crassostrea hongkongensis* | Mollusca | Bivalvia | Yes | DEGSIFEVKATAGD | |
| 4 | HSP70cA1 | JH818426.1 | EKC30019.1 | *Crassostrea gigas* | Mollusca | Bivalvia | Yes | DEGSIFEVRSTAGD | |
| 4 | | KM280636.1 | AKE47619.1 | *Ruditapes philippinarum* | Mollusca | Bivalvia | Yes | DEGSMFEVRSTAGD | |
| 4 | | KX085109.1 | AOR17366.1 | *Mizuhopecten yessoensis* | Mollusca | Bivalvia | Yes | DEGSIFEVKATAGD | |
| 4 | | KX085110.1 | AOR17367.1 | *Mizuhopecten yessoensis* | Mollusca | Bivalvia | Yes | DEGSIFEVKATAGD | |
| 4 | | KY660263.1 | AVK87132.1 | *Anadara broughtonii* | Mollusca | Bivalvia | Yes | DEGSIFEVKATAGD | |
| 4 | | XM_021507549.1 | XP_021363224.1 | *Mizuhopecten yessoensis* | Mollusca | Bivalvia | Yes | DEGSMFEVRSTAGD | |
| 4 | | XM_021514808.1 | XP_021370483.1 | *Mizuhopecten yessoensis* | Mollusca | Bivalvia | Yes | DEGSIFEVRSTAGD | |
| 4 | | XM_022444014.1 | XP_022299722.1 | *Crassostrea virginica* | Mollusca | Bivalvia | Yes | DEGSIFEVLSTAGD | |
| 4 | | XM_022459720.1 | XP_022315428.1 | *Crassostrea virginica* | Mollusca | Bivalvia | Yes | DEGSIFEVRSTAGD | |
| 4 | | XM_022459721.1 | XP_022315429.1 | *Crassostrea virginica* | Mollusca | Bivalvia | Yes | DEGSIFEVRSTAGD | |
| 4 | | XM_022462566.1 | XP_022318274.1 | *Crassostrea virginica* | Mollusca | Bivalvia | Yes | DEGSIFEVRSTAGD | |
| 4 | | XM_033886290.1 | XP_033742181.1 | *Pecten maximus* | Mollusca | Bivalvia | Yes | DEGSMFEVRSTAGD | |
| 4 | | KF195923.1 | AHB34936.1 | *Octopus vulgaris* | Mollusca | Cephalopoda | Yes | DQGSLFEVKATAGD | |
| 4 | | XM_029790319.2 | XP_029646179.2 | *Octopus sinensis* | Mollusca | Cephalopoda | Yes | DQGSLFEVKATAGD | |
| 4 | HSP70cA1 | FJ812177.1 | ACO36048.1 | *Haliotis diversicolor* | Mollusca | Gastropoda | Yes | DEGSMFEVRSTAGD | |
| 4 | | L44127.1 | AAB95297.1 | *Biomphalaria glabrata* | Mollusca | Gastropoda | Yes | DEGSIFEVKATAGD | |
| 4 | | MH220526.1 | AXR98608.1 | *Haliotis fulgens* | Mollusca | Gastropoda | Yes | DEGSMFEVRSTAGD | |
| 4 | | MK840956.1 | QKS68348.1 | *Onchidium reevesii* | Mollusca | Gastropoda | - | – | Partial sequence without the Ser-containing region; Not used included in |
| 4 | HSP70cA2 | XM_005100297.3 | XP_005100354.1 | *Aplysia californica* | Mollusca | Gastropoda | Yes | DEGSMFEVKATAGD | |
| 4 | | XM_005100299.3 | XP_005100356.1 | *Aplysia californica* | Mollusca | Gastropoda | Yes | DEGSMFEVKATAGD | |
| 4 | HSP70cA1 | XM_005103777.3 | XP_005103834.1 | *Aplysia californica* | Mollusca | Gastropoda | Yes | DEGSIFEVKATAGD | |
| 4 | HSP70cA2 | XM_009047345.1 | XP_009045593.1 | *Lottia gigantea* | Mollusca | Gastropoda | Yes | DEGSLFEVKSTAGD | |
| 4 | HSP70cA1 | XM_009053468.1 | XP_009051716.1 | *Lottia gigantea* | Mollusca | Gastropoda | Yes | DEGSLFEVKSTAGD | |
| 4 | HSP70cA3 | XM_009053469.1 | XP_009051717.1 | *Lottia gigantea* | Mollusca | Gastropoda | Yes | DEGSLFEVKSTAGD | |
| 4 | HSP70cA4 | XM_009058212.1 | XP_009056460.1 | *Lottia gigantea* | Mollusca | Gastropoda | No | E-DGVFEVLATAGD | Ser is missing as reported in Yu et al. (2021) |
| 4 | | XM_013216693.1 | XP_013072147.1 | *Biomphalaria glabrata* | Mollusca | Gastropoda | Yes | DEGSMFEVKATAGD | |

| | | | | | | | | |
|---|---|---|---|---|---|---|---|---|
| 4 | XM_013226102.1 | XP_013081556.1 | *Biomphalaria glabrata* | Mollusca | Gastropoda | Yes | DEGSMFEVKATAGD | |
| 4 | XM_013240932.1 | XP_013096386.1 | *Biomphalaria glabrata* | Mollusca | Gastropoda | Yes | DEGSMFEVKATAGD | |
| 4 | XM_025243705.1 | XP_025099490.1 | *Pomacea canaliculata* | Mollusca | Gastropoda | Yes | DEGSIFEVKATAGD | |
| 4 | XM_025245033.1 | XP_025100818.1 | *Pomacea canaliculata* | Mollusca | Gastropoda | Yes | DEGSIFEVKATAGD | |
| 4 | FN667017.1 | CBJ55211.1 | *Brissopsis lyrifera* | Echinodermata | Echinoidea | No | DD-GIFEVKSTAGD | Included in Fig. 3 |
| 4 | EU930813.1 | ACJ54702.1 | *Apostichopus japonicus* | Echinodermata | Holothuridea | No | DD-GMFEVKSTAGD | Included in Fig. 3 |
| 4 | XM_002731867.2 | XP_002731913.1 | *Saccoglossus kowalevskii* | Hemichordata | Enteropneusta | No | EDG-IFEVKSTAGD | Included in Fig. 3 |
| 4 | XM_002735377.2 | XP_002735423.1 | *Saccoglossus kowalevskii* | Hemichordata | Enteropneusta | No | EDG-IFEVKSTAGD | Included in Fig. 3 |
| 4 | XM_035626781.1 | XP_035482674.1 | *Scophthalmis maximus* | Chordata | Actinopteri | No | -EDGIFEVKSTAGD | Included in Fig. 3 |
| 4 | LR785876.1 | CAB3254511.1 | *Phallusia mammillata* | Chordata | Ascidiacea | Yes | DEGSLFEVRSTAGD | Included in Fig. 3 |
| 4 | NM_001033834.1 | NP_001029006.1 | *Ciona intestinalis* | Chordata | Ascidiacea | Yes | DEGSIFEVLSTAGD | Included in Fig. 3 |
| 4 | XM_004921180.2 | XP_004921237.1 | *Heterocephalus glaber* | Chordata | Mammalia | No | DAG-VFEVKATFGD | Included in Fig. 3 |
| 4 | XM_024116835.2 | XP_023972603.2 | *Physeter catodon* | Chordata | Mammalia | No | DA-GVFEVKATAGD | Included in Fig. 3 |
| 4 | XM_036853026.1 | XP_036708921.1 | *Balaenoptera musculus* | Chordata | Mammalia | No | D-AGVFEVKVTAGD | Included in Fig. 3 |
| 4 | XM_036922815.1 | XP_036778710.1 | *Manis pentadactyla* | Chordata | Mammalia | No | -DASVFEVKATAGD | There's a Ser but not an insersion. Included in Fig. 3 |

1 Proteins annotated by Yu et al. (2021) are underlined
