## Supplementary Table 4 for "Genomic loss of the HSP70cA gene in the vertebrate lineage"

Supplementary Table 4. HSP70s used for the phylogenetic analysis

| No. in Fig. 2 | Protein accession | Registered name | Proposed name | Scientific name | Phylum | Class | Ser |
|---|---|---|---|---|---|---|---|
| Cysotolic lineage A | | | | | | | |
| 1 | UAJ82472.1 | heat shock protein 70 kDa, partial | HSP70cA1 | *Pentalonia nigronervosa* | Arthropoda | Insecta | Yes |
| 2 | XP_025407024.1 | heat shock protein 70 A1, partial [Sipha flava] | HSP70cA1 | *Sipha flava* | Arthropoda | Insecta | Yes |
| - | BAF69068.1 | HSP70 | HSP70cA1[2] | *Bombyx mori* | Arthropoda | Insecta | Yes |
| - | NP_731651.1 | Hsp70Aa | HSP70cA1[1,2] | *Drosophila melanogaster* | Arthropoda | Insecta | Yes |
| 14 | CAB3254511.1 | hsp70 heat shock protein 70 | HSP70cA1 | *Phallusia mammillata* | Chordata | Ascidiacea | Yes |
| 15 | NP_001029006.1 | heat shock protein 70 | HSP70cA1 | *Ciona intestinalis* | Chordata | Ascidiacea | Yes |
| - | XP_029210797.1 | heat shock protein 68-like | HSP70cA1 | *Acropora millepora* | Cnidaria | Anthozoa | Yes |
| 6 | CCQ18651.1 | Molecular chaperones GRP78/BiP/KAR2, HSP70 superfamily | HSP70cA1 | *Sycon ciliatum* | Porifera | Calcerea | No |
| 10 | CBJ55211.1 | putative heat shock protein 70 | HSP70cA1 | *Brissopsis lyrifera* | Echinodermata | Echinoidea | No |
| 11 | ACJ54702.1 | heat shock protein 70 | HSP70cA1 | *Apostichopus japonicus* | Echinodermata | Holothuridea | No |
| 12 | XP_002731913.1 | PREDICTED: heat shock cognate 71 kDa protein-like | HSP70cA1 | *Saccoglossus kowalevskii* | Hemichordata | Enteropneusta | No |
| 13 | XP_002735423.1 | PREDICTED: heat shock cognate 71 kDa protein-like | HSP70cA2 | *Saccoglossus kowalevskii* | Hemichordata | Enteropneusta | No |
| 9 | CAA74243.1 | heat shock protein 70 | HSP70cA1 | *Rhabdocalyptus dawsoni* | Porifera | Hexactinellida | No |
| Cysotolic lineage B | | | | | | | |
| 17 | XP_023972603.2 | LOW QUALITY PROTEIN: heat shock 70 kDa protein 6 | HSP70cB1 | *Physeter catodon* | Chordata | Mammalia | No |
| 18 | XP_036708921.1 | LOW QUALITY PROTEIN: heat shock 70 kDa protein 6 | HSP70cB1 | *Balaenoptera musculus* | Chordata | Mammalia | No |
| 19 | XP_004921237.1 | LOW QUALITY PROTEIN: heat shock 70 kDa protein 6 | HSP70cB1 | *Heterocephalus glaber* | Chordata | Mammalia | No |
| 20 | XP_036778710.1 | LOW QUALITY PROTEIN: heat shock 70 kDa protein 6 | HSP70cB1 | *Manis pentadactyla* | Chordata | Mammalia | No |
| - | NP_005336.3 | heat shock 70 kDa protein 1A (HSPA1A) | HSP70cB1[2] | *Homo sapiens* | Chordata | Mammalia | No |
| - | NP_005518.3 | heat shock 70 kDa protein 1-like (HSPA1L) | HSP70cB3[2] | *Homo sapiens* | Chordata | Mammalia | No |
| 5 | XP_007988239.2 | LOW QUALITY PROTEIN: heat shock cognate 71 kDa protein-like | HSP70cB1 | *Chlorocebus sabaeus* | Chordata | Mammalia | Yes |
| - | P19120.2 | Heat shock cognate 71 kDa protein | HSP70cB1[2] | *Bos taurus* | Chordata | Mammalia | No |
| - | NP_077327.1 | heat shock cognate 71 kDa protein | HSP70cB1[2] | *Rattus norvegicus* | Chordata | Mammalia | No |
| - | AAB21658.1 | HSC71 | HSP70cB1[2] | *Oncorhynchus mykiss* | Chordata | Actinopteri | No |
| 16 | XP_035482674.1 | heat shock cognate 71 kDa protein-like | HSP70cB1 | *Scophthalmus maximus* | Chordata | Actinopteri | No |
| - | BAF94142.1 | heat shock protein 70A | HSP70cB1[2] | *Alligator mississippiensis* | Chordata | Sarcopterygii (superclass) | No |
| - | CAH04109.1 | heat shock cognate 71 | HSP70cB2[2] | *Mytilus galloprovincialis* | Mollusca | Bivalvia | No |
| 7 | XP_011404198.1 | heat shock protein 70 B2 | HSP70cB2[2] | *Amphimedon queenslandica* | Porifera | Demospongiae | No |
| 8 | XP_011404208.1 | PREDICTED: heat shock cognate 71 kDa protein-like | HSP70cB1[2] | *Amphimedon queenslandica* | Porifera | Demospongiae | No |
| - | XP_029206535.1 | heat shock cognate 71 kDa protein-like | HSP70cB1[2] | *Acropora millepora* | Cnidaria | Anthozoa | No |
| - | XP_029194855.1 | heat shock cognate 71 kDa protein-like | HSP70cB2[2] | *Acropora millepora* | Cnidaria | Anthozoa | No |
| 3 | XP_033624415.1 | heat shock cognate 71 kDa protein-like | HSP70cB1 | *Asterias rubens* | Echinodermata | Asteroidea | No |
| 4 | XP_033625279.1 | heat shock cognate 71 kDa protein | HSP70cB2 | *Asterias rubens* | Echinodermata | Asteroidea | No |
| - | XP_014783074.1 | PREDICTED: heat shock cognate 71 kDa protein | HSP70cB1[2] | *Octopus bimaculoides* | Mollusca | Cephalopoda | No |

| | | | | | | | | |
|---|---|---|---|---|---|---|---|---|
| | - | NP_524063.1 | heat shock protein cognate 1, isoform A | HSP70cB1[1] | *Drosophila melanogaster* | Arthropoda | Insecta | No |
| | - | NP_524356.1 | heat shock protein cognate 4, isoform A | HSP70cB2[1] | *Drosophila melanogaster* | Arthropoda | Insecta | No |
| Endoplasmic reticulum | | | | | | | | |
| | - | XP_029187601.1 | endoplasmic reticulum chaperone BiP-like | HSP70er1[2] | *Acropora millepora* | Cnidaria | Anthozoa | - |
| | - | NP_005338.1 | endoplasmic reticulum chaperone BiP precursor | HSP70er1[2] | *Homo sapiens* | Chordata | Mammalia | - |
| | - | NP_727563.1 | heat shock 70-kDa protein cognate 3, isoform A | HSP70er1[1] | *Drosophila melanogaster* | Arthropoda | Insecta | - |
| Mitochondria | | | | | | | | |
| | - | XP_029208733.1 | stress-70 protein, mitochondrial-like | HSP70m1[2] | *Acropora millepora* | Cnidaria | Anthozoa | - |
| | - | NP_004125.3 | stress-70 protein, mitochondrial precursor (HSPA9) | HSP70m1[2] | *Homo sapiens* | Chordata | Mammalia | - |
| | - | NP_523741.2 | heat shock protein cognate 5, isoform A | HSP70m1[1] | *Drosophila melanogaster* | Arthropoda | Insecta | - |
| Yeast | | | | | | | | |
| | - | YAL005C | SSA1 (Cyt) | - | *Saccharomyces cerevisiae* | Ascomycota | Saccharomycetes | No |
| | - | YLL024C | SSA2 (Cyt) | - | *Saccharomyces cerevisiae* | Ascomycota | Saccharomycetes | No |
| | - | YBL075C | SSA3 (Cyt) | - | *Saccharomyces cerevisiae* | Ascomycota | Saccharomycetes | No |
| | - | YER103W | SSA4 (Cyt) | - | *Saccharomyces cerevisiae* | Ascomycota | Saccharomycetes | No |
| | - | YJL034W | KAR2 (ER) | - | *Saccharomyces cerevisiae* | Ascomycota | Saccharomycetes | No |
| | - | YJR045C | SSC1 (Mt) | - | *Saccharomyces cerevisiae* | Ascomycota | Saccharomycetes | No |

[1]Submitted microPublication Biology

[2]Yu et al. (2021)

3Grawal et al. (2022)
